# Exploring biological networks in 3D, stereoscopic 3D and immersive 3D with iCAVE

**DOI:** 10.1101/212126

**Authors:** Selim Kalaycı, Zeynep H. Gümüş

## Abstract

Biological networks are becoming increasingly large and complex, pushing the limits of existing 2D tools. iCAVE is an open source software tool for interactive visual explorations of large and complex networks in 3D, stereoscopic 3D or immersive 3D. It introduces new 3D network layout algorithms and 3D-extensions of popular 2D network layout, clustering and edge bundling algorithms to assists researchers in understanding the underlying patterns in large, multi-layered, clustered or complex networks. This protocol aims to guide new users on the basic functions of iCAVE for loading data, laying out networks (single or multi-layered), bundling edges, clustering networks, visualizing clusters, visualizing data attributes and saving output images or videos. It also provides examples on visualizing networks constrained in physical 3D space (e.g. proteins; neurons; brain). It is accompanied with a new version of iCAVE with an enhanced user interface and highlights new features useful for existing users.

**Significance Statement:** Network representations assist in systems-level data exploration in many research fields, providing valuable insights. However, with the recent advances in experimental technologies, biological networks are becoming increasingly large and complex, necessitating new data visualization solutions. We have recently developed iCAVE (Liluashvili et al., 2017), an open?source software platform that enables 3D (optionally stereoscopic and or immersive) visualizations of complex, dense or multi?layered biological networks. Users can select from several new 3D network layout and clustering algorithms, bundle network edges and customize network attributes to reveal hidden structures within them. This protocol guides new users on loading, navigating, customizing and saving networks in iCAVE and is accompanied by an updated version of the software.

## INTRODUCTION

Networks are frequently used to visually communicate and understand complex associations in biomedical data (Merico, Gfeller, & Bader, 2009; M. Newman, 2010). There are myriad tools that assist in interactively exploring networks in two dimensions (2D) including Cytoscape (Shannon et al., 2003), Gephi (Bastian, Heymann, & Jacomy, 2009), and Osprey (Breitkreutz, Stark, & Tyers, 2003). However, recent developments in both computational and experimental technologies are leading to increasingly large and heterogeneous data with ever more complex interactions, pushing the boundaries of existing 2D tools, which are inherently limited by the number of interactions that can fit within a 2D screen. New visualization approaches are needed to address the challenges in exploring such datasets (Pavlopoulos et al., 2015). To address these needs, we have recently introduced iCAVE, an open-source software tool for visualizations of complex, large and/or multi-layered networks in 3D, stereoscopic 3D and immersive 3D (Liluashvili et al., 2017). For networks that represent abstract data, the addition of the third dimension allows for greater flexibility, especially when combined with rotation, zoom and navigation capabilities (Sollenberger & Milgram, 1993). For networks where node coordinates have physical meaning within the 3D space (e.g. protein residue-residue or brain connectivity networks), users can navigate within the actual physical layout of their networks.

This protocol aims to guide new users to gain hands-on experience in using the basic functions of iCAVE to interactively explore networks in 3D. Users that follow the protocols will learn loading data, laying out networks (single or multi-layered), bundling edges, clustering networks, visualizing clusters, visualizing different data attributes (e.g. colors, sizes, directions) and saving iCAVE outputs of network statistics reports, image snapshots and animation videos. The protocols are illustrated with multiple examples and 2D snapshots of their output images in the accompanying figures. Their input files are provided in the Internet Resources. For a couple networks, we also provide their videos. These networks can then be easily shared with colleagues or reproduced.

Briefly, Basic Protocol 1 describes how to customize, load and navigate network data and to run different network layout algorithms within iCAVE. Basic Protocol 2 details how to visualize and explore interconnections in multi-layered networks and how to bundle edges in order to reveal connectivity strengths between different data types. Basic Protocol 3 provides information on running different network-topology based graph clustering algorithms, cluster visualization options and additional utilizations of edge bundling to reveal intra- and inter-cluster connectivity strengths. Finally, Basic Protocol 4 details how to visualize networks with known 3D coordinates. In addition, Supplementary Protocol 1 provides instructions on installing iCAVE and Supplementary Protocol 2 details both the steps to turn stereo option on or off for stereoscopic 3D viewing, and the necessary hardware and software resources. Finally, this manuscript is further accompanied with a new version of iCAVE that has an enhanced user interface, and its new features are summarized in the Background section. Overall, this protocol will assist researchers in a variety of fields to visually explore complex networks in 3D and to identify and communicate special structures within them using iCAVE.

## BASIC PROTOCOL 1

### VISUALIZING NETWORK DATA FROM INPUT FILE IN iCAVE

Users can download iCAVE from http://research.mssm.edu/gumuslab/software.html and follow the installation instructions provided in Support Protocol 1. After successful installation, all examples provided in this protocol can be generated in 3D using iCAVE. For additional stereo 3D viewing, necessary resources and details on turning stereo on or off are provided in Support Protocol 2.

In Basic Protocol I, we demonstrate how to customize, load, layout, navigate and save statistics and video animations of a network from (Goh et al., 2007) on disease-disease and disease-gene associations in a *diseasome* network (see “diseasome.txt” in Internet Resources). We use the most common iCAVE input, which is a a tab-delimited text file of all edges that is customized to user specifications, though it is also possible to input only a list of gene names for a quick, initial query of iCAVE's COMBO database (see (Liluashvili et al., 2017) for details of this). The input data is passed as an argument to iCAVE during execution time, based on which it generates an interactive visualization of the network in 3D space. Users can select from any of its layout algorithms, which include both novel 3D algorithms and extensions of popular 2D algorithms to 3D. The network layout step is important as it can affect understandings of its structure. Users are encouraged to test all layout algorithms within iCAVE to find the most suitable for their networks. It will be different for each network. They can then navigate within the network by interactively zooming and rotating, view network statistics or save network animations at user defined speed, zoom or viewpoints to communicate the network with colleagues.

#### Necessary Resources

##### Input File Format

In general, iCAVE input is a tab-separated text file with two sections: an optional section (at the top of the file) and a required section (below). The optional section lists the nodes and the required section lists the node pairs that make up the edges.

The optional section begins with the header:

~~~
#nodes name
~~~

The required section starts immediately below (no spaces) with:

~~~
#edges node1 node2
~~~

Both sections may also include user-defined attributes for nodes and edges. All attribute options are summarized in Support Protocol 3. For example, in input file “diseasome.txt”, which includes 1,419 nodes and 2,738 edges, the optional nodes section begins with:

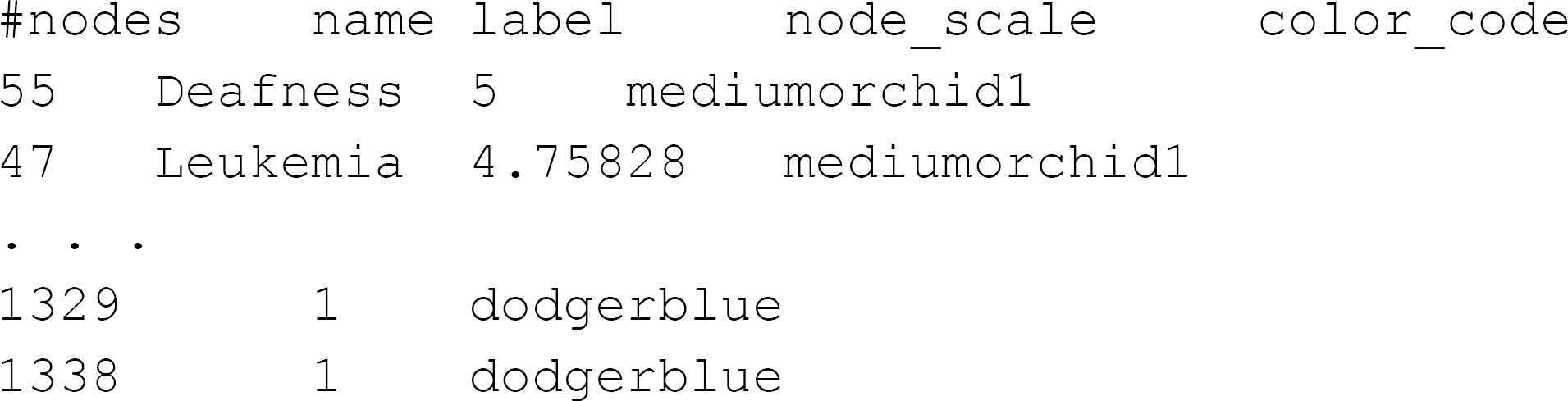

Here, the header line includes the required node ‘name’ attribute and the optional ‘label’, ‘node_scale’, and ‘color_code’ attributes. Each node has a unique name. Additional user-defined labels are specified under the optional ‘label’ attribute for all or a subset of nodes. The optional ‘node_scale’ attribute enables node size scaling based on user-defined criteria. Finally, the optional ‘color_code’ attribute enables user-defined color specifications for every node. In this example, the two lines below the header line specify nodes that represent diseases, and the last two lines specify nodes that represent genes. Disease nodes have descriptive labels which we will interactively turn on or off during this session, while labels for gene nodes are left blank. Diseases are ranked in decreasing order of their user-defined node_scale values (starting from 5) that specify the disease node size based on their scale, while all gene nodes are assigned a node_scale value of 1. Here, to differentiate between node types, all disease nodes are user-defined in color ‘mediumorchid1’, and all gene nodes in ‘dodgerblue’.

In diseasome network, the first few lines of the required edges section are:

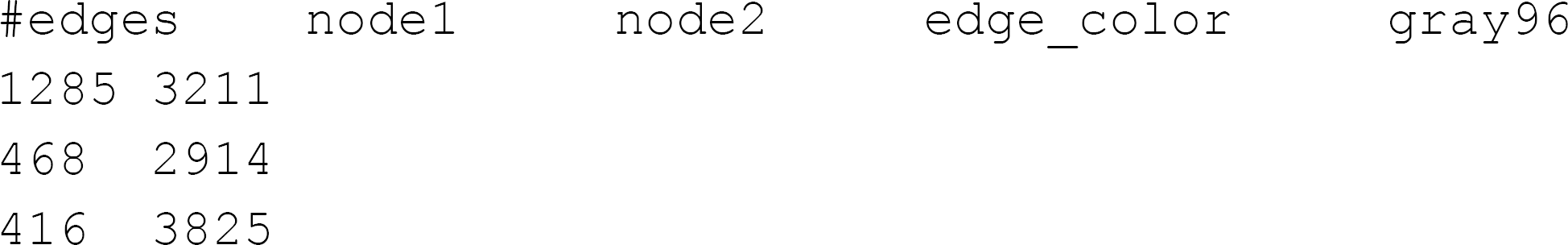

The header includes the required ‘node1’ and ‘node2’ attributes and an optional ‘edge_color’ attribute that assigns the same color to all edges (alternatively, using the ‘color_code’ option, users can define color for each edge separately). Each line below the header corresponds to an individual edge. In this example, the first edge is specified between nodes 1285 and 3211, and the color for all edges is specified as ‘gray96’.

Protocol steps – *step annotations*

1. Copy the input file “diseasome.txt” into $iCAVE_HOME/vrnet/importFiles folder.
2. Open a terminal and set your directory to $iCAVE_HOME/vrnet.
3. Launch iCAVE with the following syntax:

~~~
vrnetview -Gdiseasome.txt -O10 -l1 -B1
~~~ `-Gdiseasome.txt` to open the graph specified in file diseasome.txt. `-010` to specify that the file is in iCAVE format. `-l1` to use weighted-force directed algorithm for network layout. Other layout options are: `-l0` = Force-directed, `-l2` = Hemispherical, `-l3` = Semantic Levels, `-l4` = LinLog, `-l5` = Weighted LinLog, `-l6` = Hybrid Force-directed (default layout if no layout option is provided), `-l7` = Simulated Annealing Force-directed, `-l8` = Coarsened Force-directed `-B1` to use white background. If it is omitted, the default black background is used.

#### Navigating the Network

4. Zoom in/out by either (i) pressing left mouse button + "z" key in the keyboard and moving the mouse, or (ii) using the mouse rotation wheel.
5. Rotate the network by pressing the left mouse button and moving the mouse. To move the whole network to your desired location in the window, press *“z”* key in the keyboard and move the mouse.
6. If no problem occurs, you should get a view similar to one as in Fig. 1. (Please note that all the images are captured via basic screenshot operations.)

**Figure 1.**
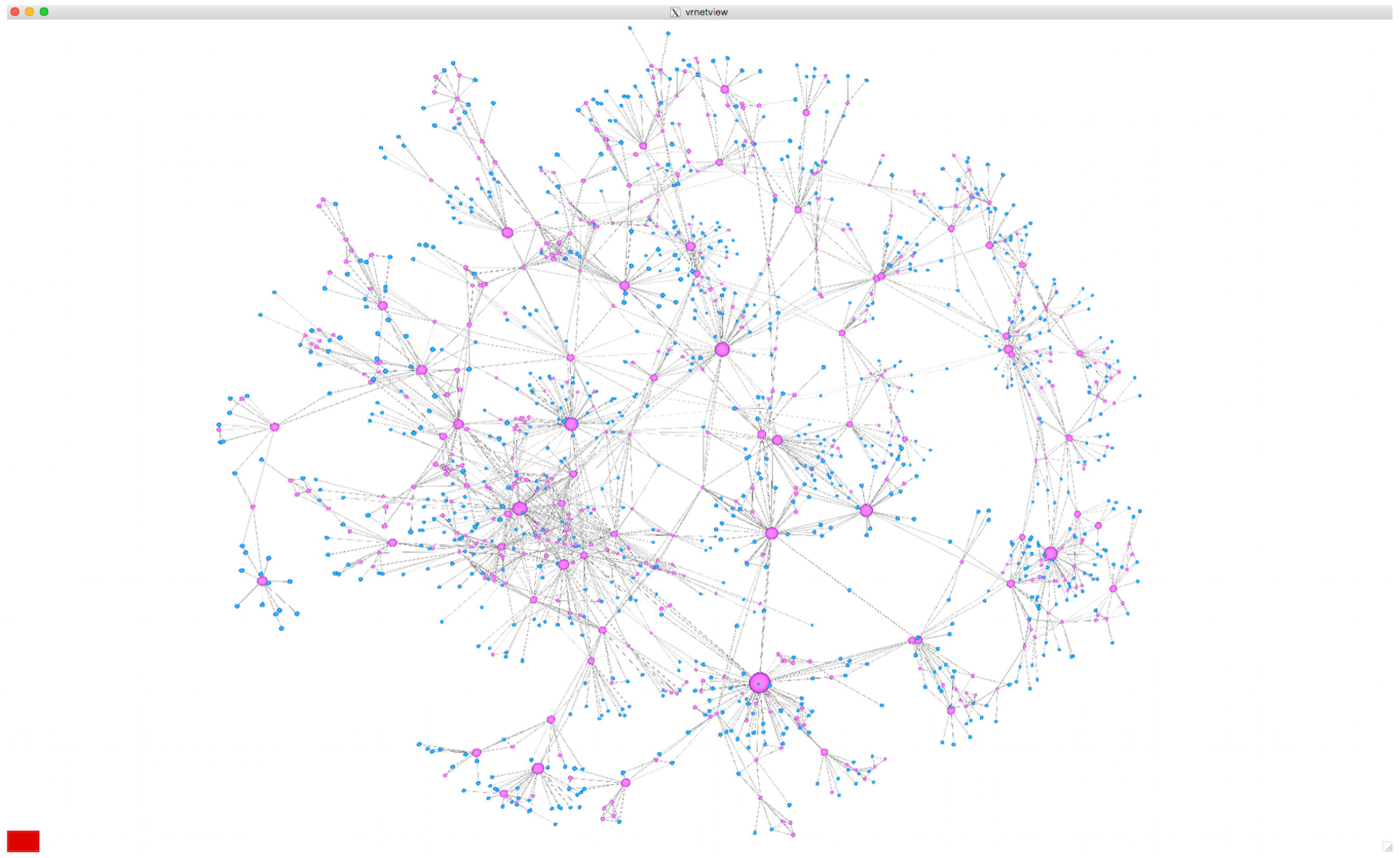
Global view of the diseasome network.

#### Network Layout Options

The current version of iCAVE includes eight different network layout algorithms. Please test each algorithm to see which works best in communicating your network. In Fig. 2, we illustrate different network layouts for the diseasome network using four of these algorithms.

**Figure 2.**
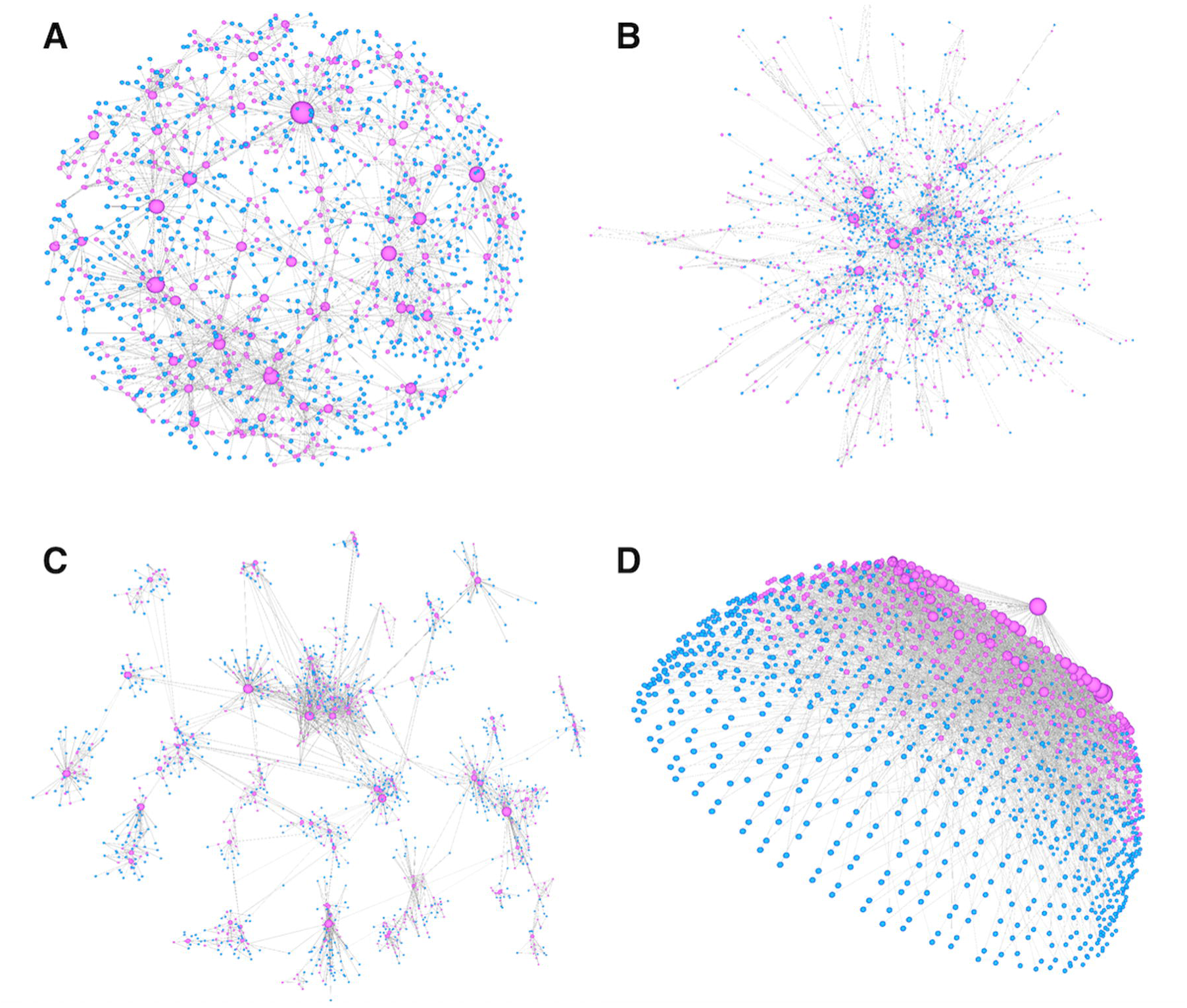
Diseasome network visualized using different layout algorithms in iCAVE. (a) Forcedirected, (b) coarsened force-directed, (c) simulated annealing force-directed, (d) hemisphere layout algorithms.

7. Open the main menu of iCAVE by pressing the right mouse button, and scroll inside “Layout Algorithms”. All available layout algorithm options are listed as toggle buttons.

i. For Fig. 2.a, select “Force Directed” toggle button.
ii. For Fig. 2.b, select “Coarsened Force Directed” toggle button.
iii. For Fig. 2.c, select “Simulated Annealing Force Directed” toggle button.
iv. For Fig. 2.d, select “Hemisphere” toggle button.
8. After testing several available layouts in iCAVE (see Fig. 2. a-d) for this network, we choose the weighted force directed layout as the most informative, as shown in Fig. 1. Similarly, users are encouraged to try different layouts to see which one works best for their particular network. In this Protocol, we will continue performing further operations on this layout. To switch to the weighted force directed layout, go to the main menu and select “Layout Algorithms —> Weighted Force Directed” toggle button. You should get a view similar to Fig. 1.

#### Adjusting Size of Network Nodes/Edges

9. To adjust node or edge size, go to main menu and select “Adjust Node/Edge Size”.
10. You should get a graphical slider as shown in Fig. 3.

**Figure 3.**
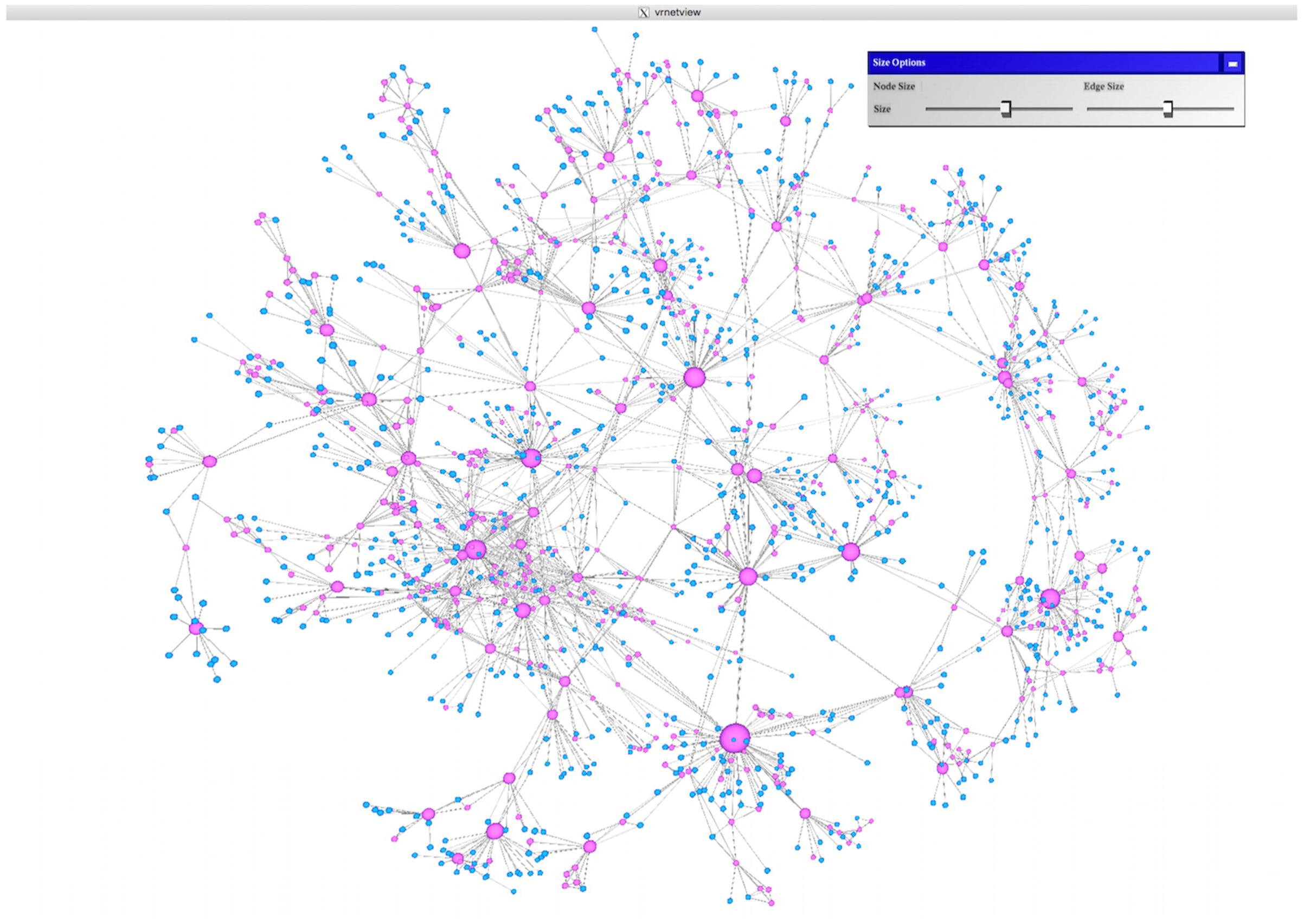
Graphical sliders to adjust the size of nodes and edges.
11. Adjust the sizes of nodes and/or edges by using the sliders.
12. Close the “Size Options” popup menu window by clicking on the close button on top-right.

#### Displaying Node Labels

13. From the main iCAVE menu, select “Show Labels”.
14. To interactively adjust the size of labels, go to main menu, select “Adjust Label Size”. Use the graphical slider to increase or decrease the size of labels.
15. Since there are a large number of nodes, you may not be able to read all network labels. For a better viewing, zoom into an area in which you are interested; now you can read all the labels. For example, you can see the region of the network comprised of diseases like Dementia, Alzheimer, and Schizophrenia in the zoomed-in area of the network in Fig 4.

**Figure 4.**
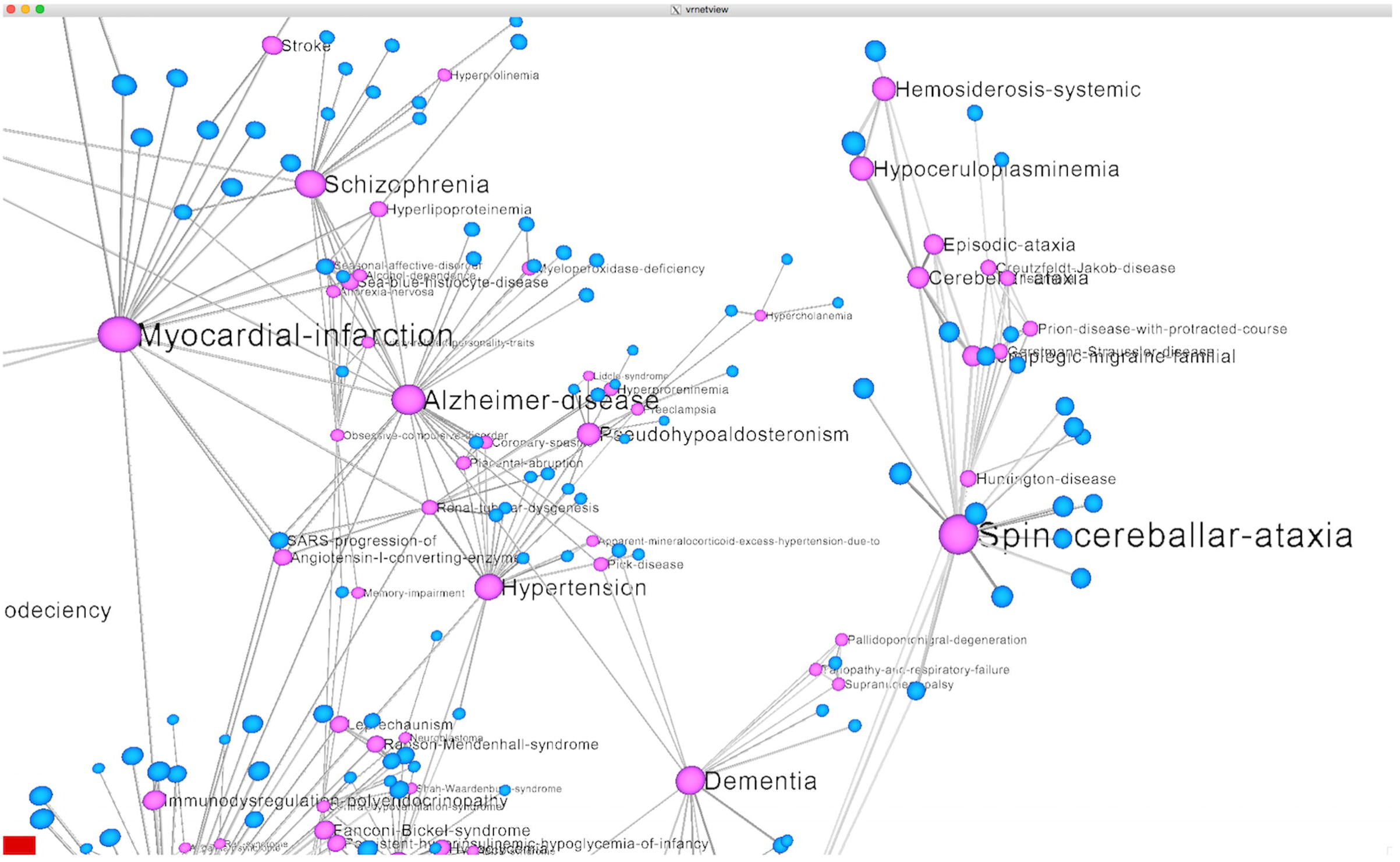
Disease labels displayed in a zoomed-in section of the diseasome network.
16. (optional) Alternatively, you can modify the input file to only display labels for some nodes. In a large network such as this one, this will draw attention to certain nodes in the global network view. As an illustration, the input file (“diseasome-maindiseases.txt”, see Internet Resources) provides node labels only for the top 13 diseases in the network.

i. Copy the input file “diseasome-maindiseases.txt” into $iCAVE_HOME/vrnet/importFiles.
ii. Open another terminal window and change your directory to $iCAVE_HOME/vrnet.
iii. Launch iCAVE by using the following in the command line:

~~~
vrnetview -Gdiseasome-maindiseases.txt -O10 -l1 -B1
~~~
iv. Follow steps 13-14 to display and adjust the labels in the network.
v. You should get a view similar to Fig. 5.

**Figure 5.**
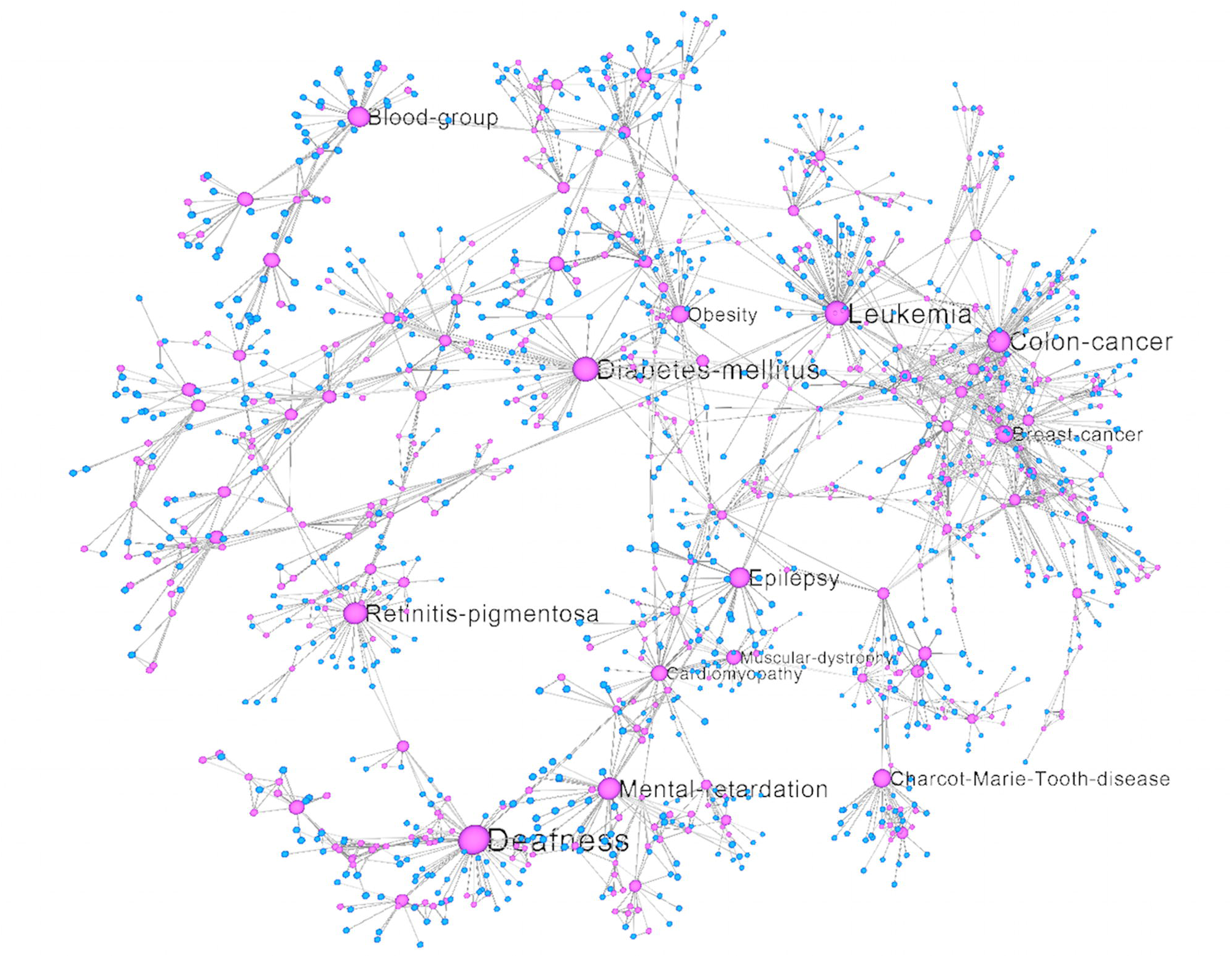
Labels for top diseases displayed in a complete view of the diseasome network.
17. You can turn node labels off to view the network structure with less occlusions. For this purpose, open the main iCAVE menu, and uncheck the “Show Labels” toggle button. You should get a global network view similar to Fig. 1.

#### Network General Measures

You can use iCAVE to calculate general network measures as summarized in Fig. 6.

**Figure 6.**
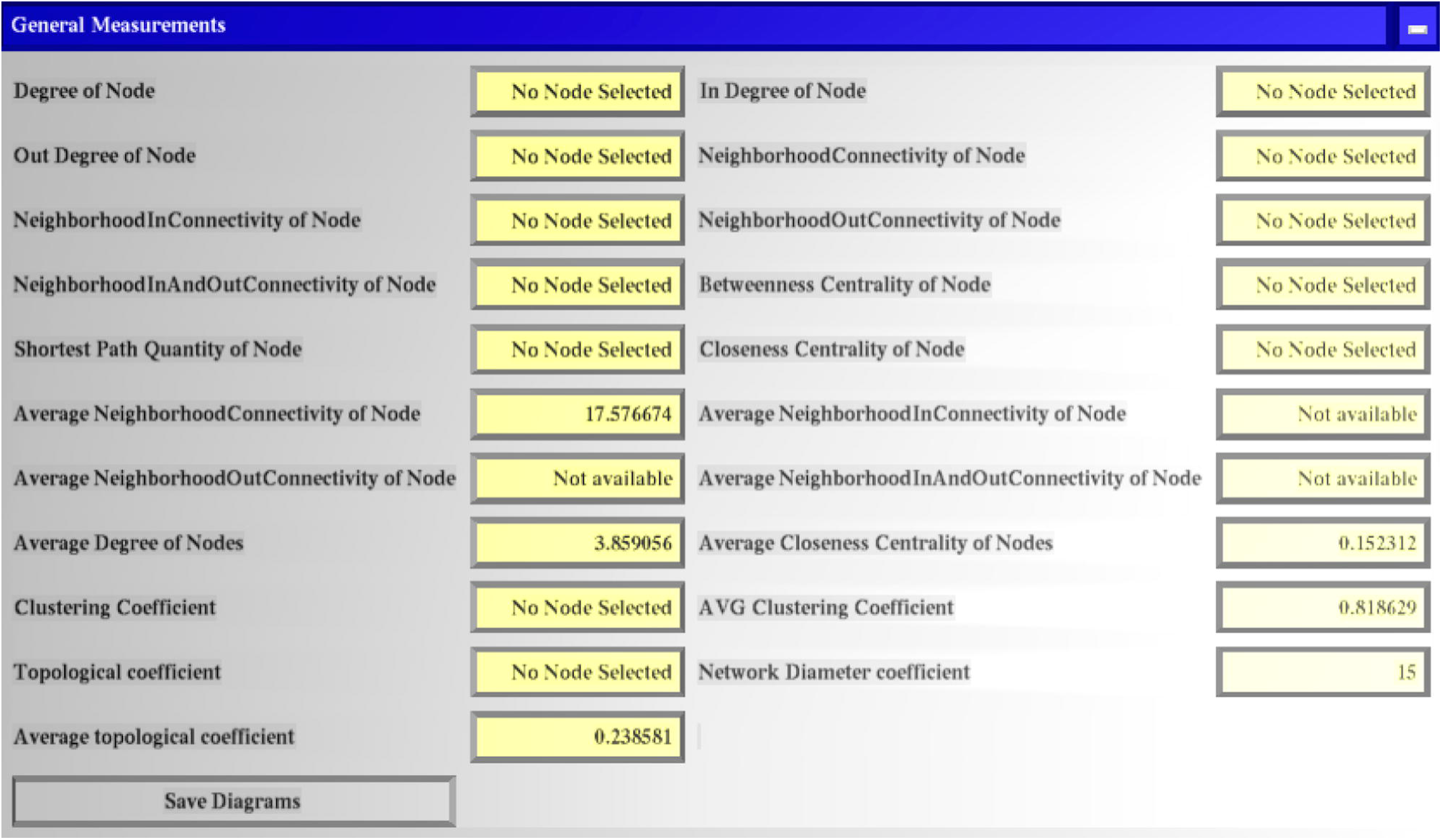
Pop-up window displaying general network measures.

18. From the main iCAVE menu, select and enable “Show General Measures” toggle.
19. You should get a view similar to Fig. 6. If you want to save the measures displayed in the window, click “Save Diagrams” option, and several text and graph files on different network measures will be saved in your “$iCAVE_HOME/vrnet/diagrams” folder.

#### Saving Animations

Using 2D snapshots of a network is not ideal, as it projects the 3D network to a lower dimension, decreasing its communication efficiency. Therefore, iCAVE provides functionality to generate and save 3D animation videos for a rotating model of the network. Users can save and share these animations to communicate their networks with others:

20. From the main iCAVE menu, select and enable “Rotate Model” toggle.
21. Use the graphical sliders in the dialog box to adjust rotation speed and direction.
22. Click the “Save Rotation as Movie” button. Animation file (see Supplementary Video 1) will be saved in your “$iCAVE_HOME/vrnet” folder and named “animation.gif” by default.

## BASIC PROTOCOL 2

### VISUALIZING MULTIPLE LAYERS OF INFORMATION IN iCAVE

Systems-level studies often pool together multiple types of information in efforts to understand their interactions within the integrated system (Butcher, Berg, & Kunkel, 2004). For example, the diseasome network discussed in Protocol 1 is a good case study for exploring multiple layers of information, as it includes two different but inter-related information types: diseases and genes. In Basic Protocol 2, we guide users on how to visualize and explore the interconnections within networks that contain multiple layers of information, using this diseasome network and demonstrate how it can be easily specified in two layers in the input file. We then further introduce edge bundling operation, which clarifies the underlying network topology by bundling adjacent edges together. Edge bundling is advantageous when a large number of connections between network layers exist, confusing the viewer. iCAVE has several user-adjustable parameters to adjust the desired degree of granularity of the bundled edges, as summarized below.

#### Necessary Resources

##### Input File

As the input file, we will use a slightly modified version of the file used in Protocol 1. This time, the disease nodes and gene nodes are specified in two separate layers (see “diseasome-layers.txt”, in Internet Resources). iCAVE will take the layer information into account in the visualization. Here are a few lines in the optional nodes section of the input file:

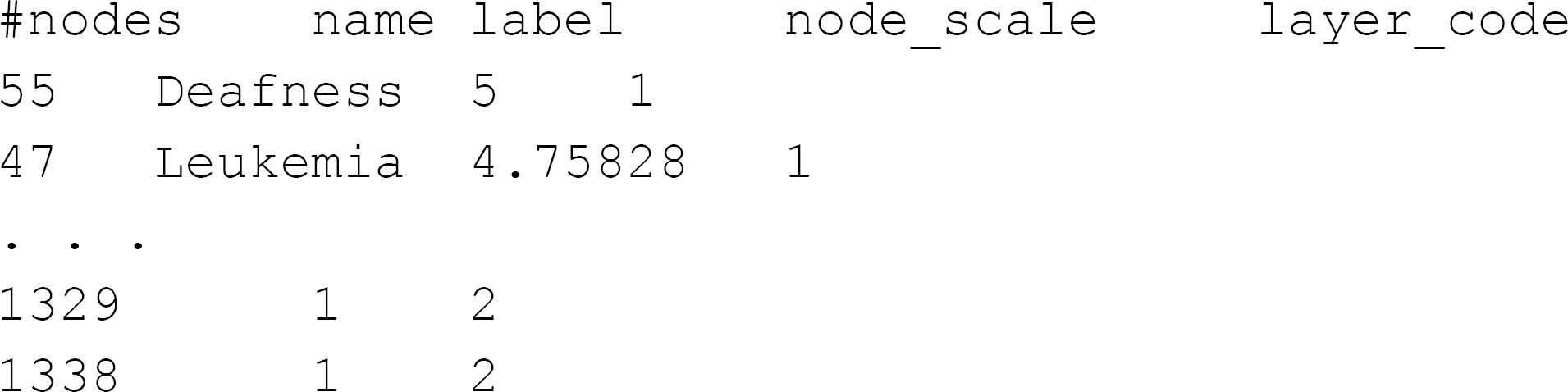

As you can see, we have altered two attributes in the nodes section. First, we removed the attribute ‘color_code’, as node colors in layers are automatically assigned by iCAVE, depending on the number of layers. Second, we added the attribute ‘layer_code”, where we specified all diseases in layer ‘1’, and all genes in layer ‘2’. The remaining information in the nodes section is the same as in Protocol 1.

Edges section is also similar to the input file in Protocol 1. The only change is on edge coloring. In this version, we will display the colors for disease-gene edges in ‘lightskyblue’, and disease-disease edges in ‘violet’. The remaining information in the edges section is the same as in Protocol 1. Here are a few lines from the top and bottom of the edges section:

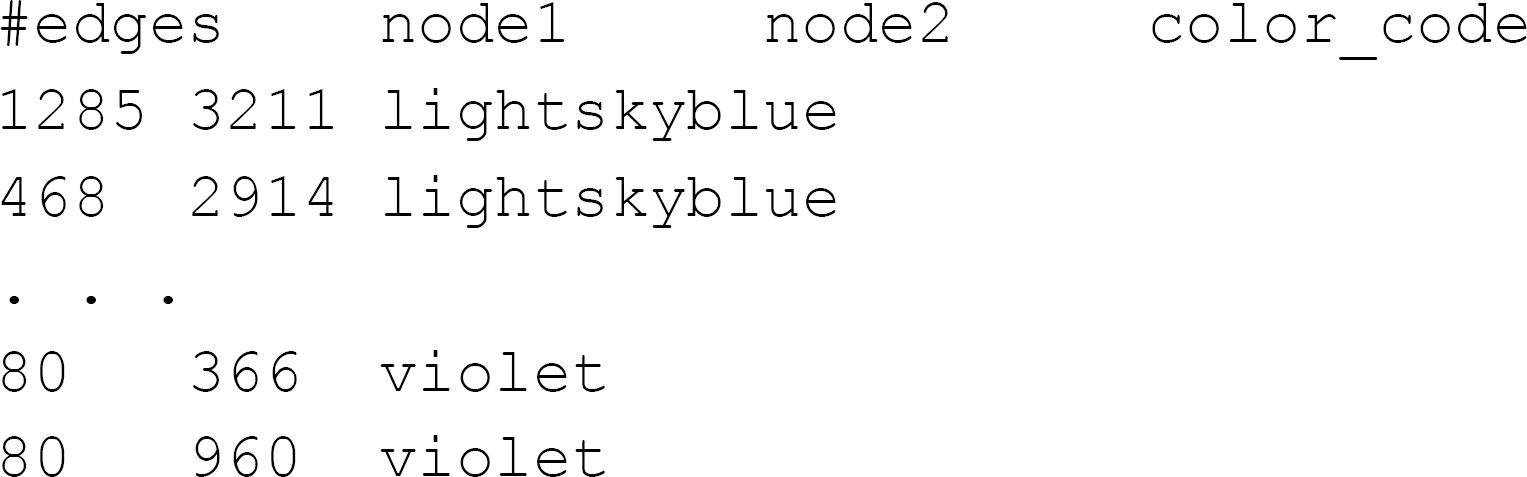

Protocol steps – *step annotations*

1. Copy the input file “diseasome-layers.txt” into $iCAVE_HOME/vrnet/importFiles folder.
2. Open a terminal and change your directory to $iCAVE_HOME/vrnet.
3. Launch iCAVE with the following syntax:

~~~
vrnetview -Gdiseasome-layers.txt -O10 -B1
~~~ `-Gdiseasome-layers.txt` to open the graph specified in file diseasome-layers.txt. `-O10` the file is in iCAVE format. `-B1` to use white color background.
4. You should get a view similar to Fig. 7. The nodes at the top layer represent diseases, and those at the bottom layer represent genes.

**Figure 7.**
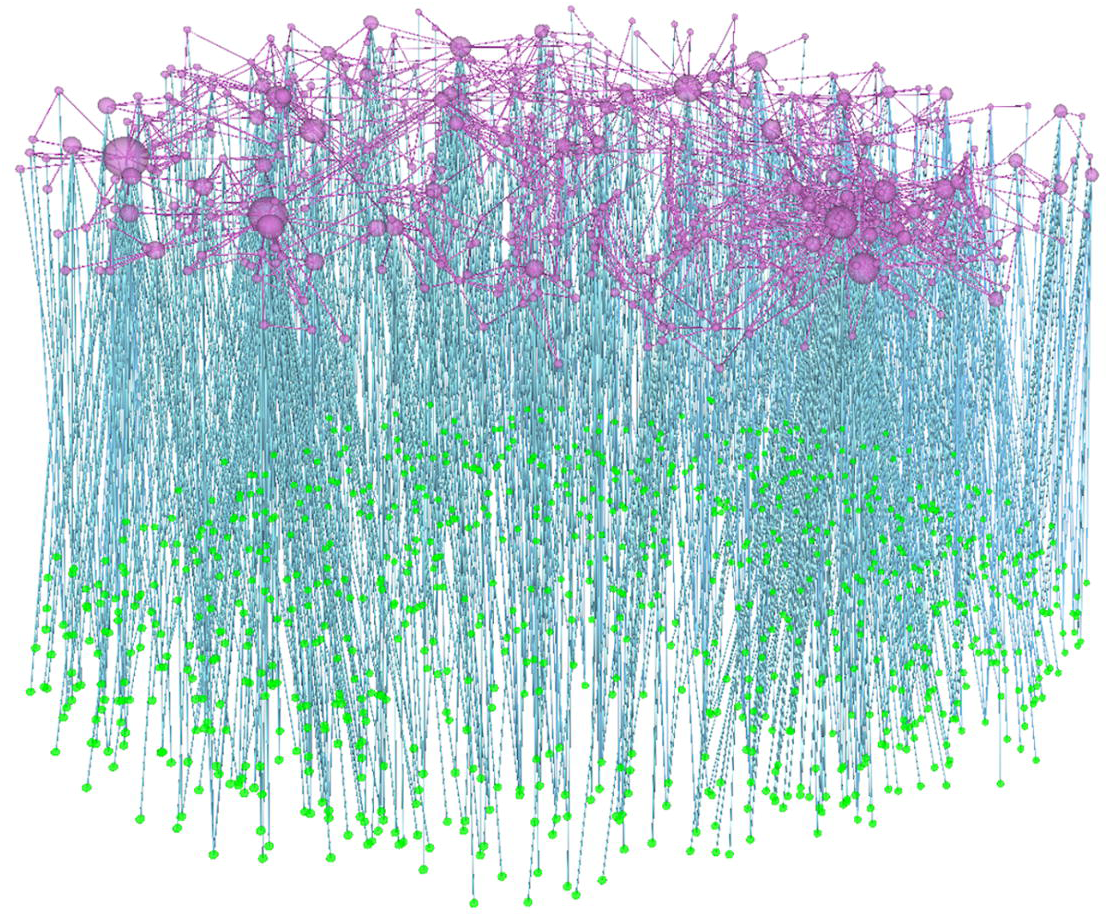
Layered view of the diseasome network.

#### Edge Bundling

Due to large number of connections between layers, it is difficult to discern topological properties of the network in Fig. 7. Edge bundling operation (Holten & van Wijk, 2009) in iCAVE visually bundles adjacent edges together, analogous to bundling electrical wires or cables. Bundling is extremely useful in identifying global patterns in very large networks. In layered networks, bundling often leads to clusters of connections between two layers based on the connectivity patterns between the two layers. Furthermore, users can alter the edge bundling parameter for finer or coarser clusters. Bundling the edges of the network in Fig. 7 enables the representation of highest-density “correlation highways” that connect diseases and genes.

To perform optional edge bundling operation:

5. From the main menu of iCAVE, select “Network Algorithms —> Bundle Edges” button.
6. In the dialog box, choose the settings as shown in Fig. 8.

**Figure 8.**
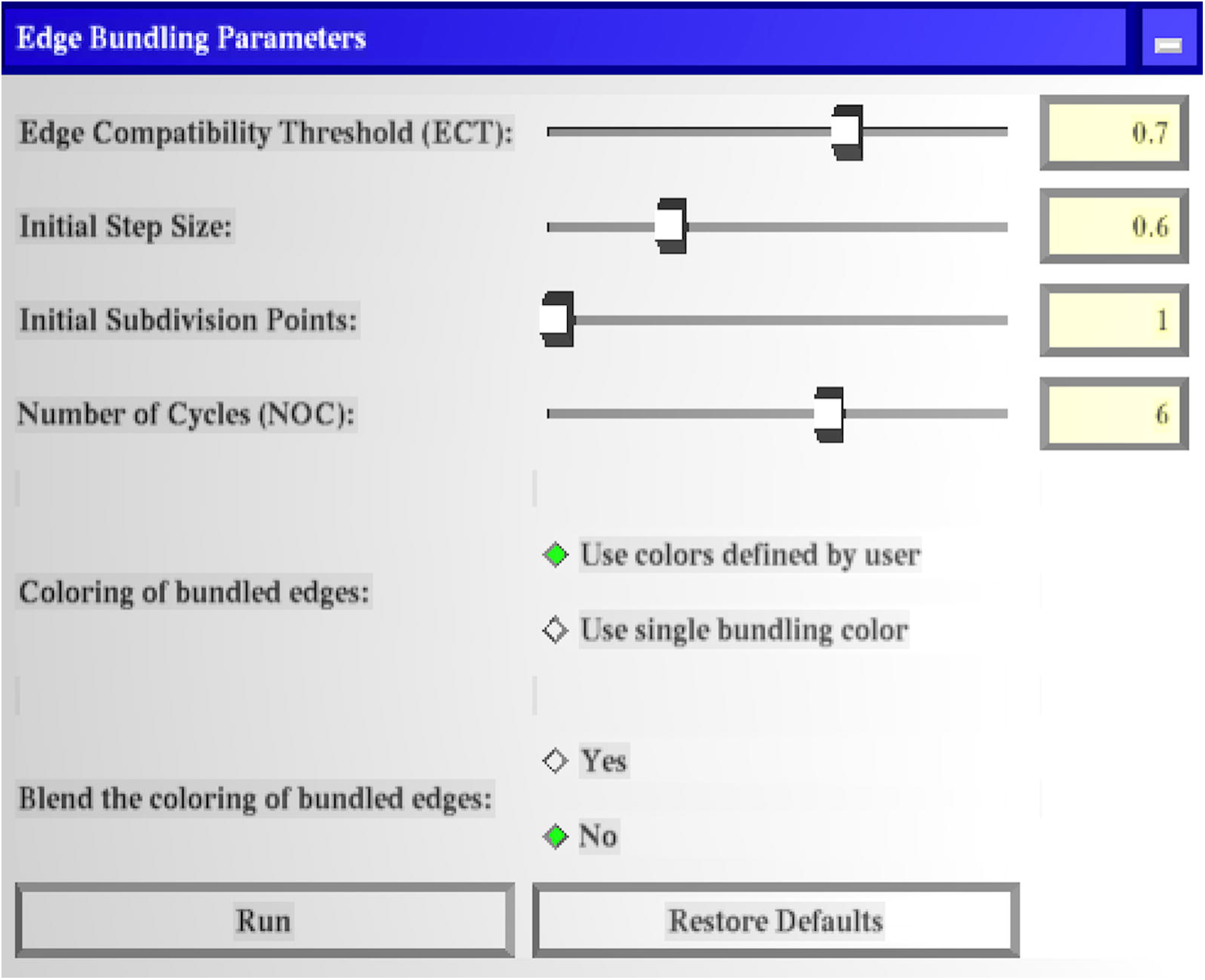
Edge bundling dialog box.
7. Click the Run button.
8. You should get a view similar to Fig. 9. Edge bundling provides a much cleaner image and more insights regarding the global topology of the network as compared to Fig. 7. Correlation highways between the diseases and genes can be explored further. Also, edge bundling can be repeated with different Edge Compatibility Threshold (ECT) values to get correlation highways at varying degrees of granularity- from very fine to very coarse clusters.

**Figure 9.**
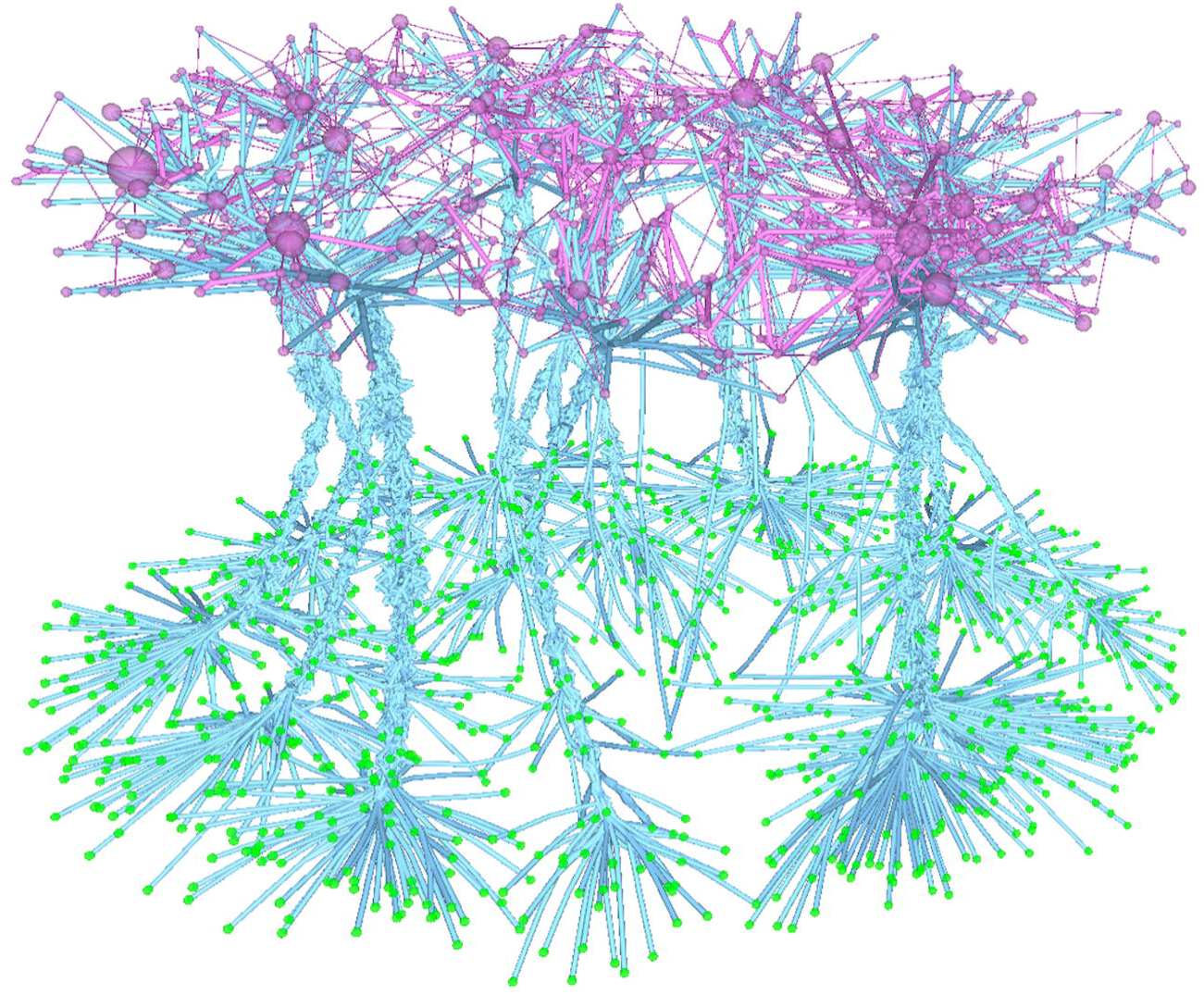
Edge bundled view of the network segregates the connections between two layers into 7 clusters or ‘highways’ of interaction. Users can alter the granularity of these connections with the ECT option.
9. To display the network in full/original view, from the main menu of iCAVE, select “Reset Options —> Reset Network” button. You should get the view in Fig. 7.

## BASIC PROTOCOL 3

### NETWORK TOPOLOGY BASED GRAPH CLUSTERING AND CLUSTER VISUALIZATIONS IN iCAVE

In Basic Protocol 3, we demonstrate network topology-based clustering algorithms and cluster visualization options provided in iCAVE, integrated with additional edge bundling. Clustering is important in understanding the network structure, and iCAVE currently offers three topology-based clustering algorithms. These are edge-betweenness (Girvan & Newman, 2002), Markov (van Dongen, 2000), and modularity (M. E. J. Newman, 2006). In addition, iCAVE provides several novel algorithms for the layout of these clusters in the 3D space. By default, iCAVE uses a force-directed algorithm for cluster layout (details in (Liluashvili et al., 2017)). Alternatively, users can layout clusters using 3D Circos option or display them in-position, where network layout stays the same but nodes are colored based on the graph clustering results. Each algorithm has strengths and weaknesses that depend on the network under study. We recommend that users test and choose the best clustering and cluster visualization method for their network, and then add edge bundling to clarify the strength of intra-cluster and inter-cluster connectivities.

#### Necessary Resources

##### Input File

We will illustrate Protocol 3 using the input file “visual-cortex.txt” (see Internet Resources), with data from (Rossi & Ahmed, 2015). This file specifies the network structure of the visual cortex of a mouse brain. There are 193 nodes and 214 edges. The top lines of the nodes section of the input file are:

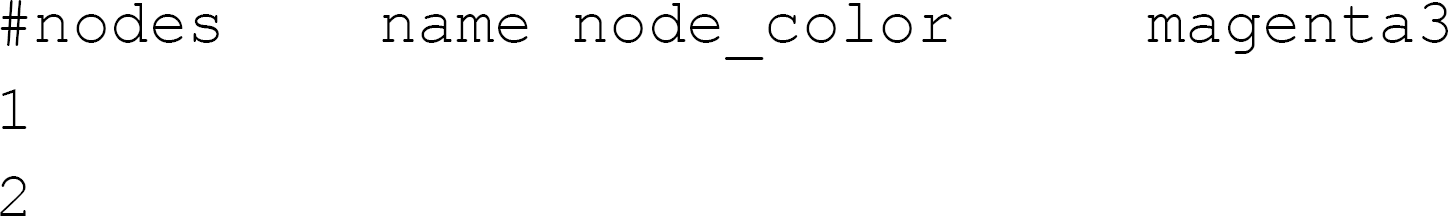

The header line includes the required ‘name’ attribute and the optional the ‘node_color’ attribute, which lets users specify a color to be used for all network nodes. The next two lines after the header list node names.

The nodes section is followed by the required edges section, which specifies all the edges in the network. First few lines of the edges section is as follows:

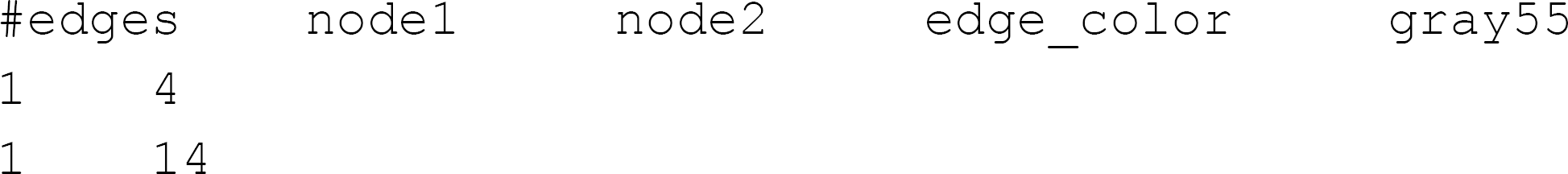

The header line includes the required attributes ‘node1’ and ‘node2’ and the optional ‘edge_color’ attribute. Each line after the header specifies an edge.

Protocol steps – *step annotations*

1. Copy the input file “visual-cortex.txt” into $iCAVE_HOME/vrnet/importFiles folder.
2. Open a terminal and change your directory to $iCAVE_HOME/vrnet.
3. Launch iCAVE with the following syntax:

~~~
vrnetview -Gvisual-cortex.txt -O10 -l1 -B1
~~~ `-Gvisual-cortex.txt` to open the graph specified in file visual-cortex.txt. `-O10` the file is in iCAVE format. `-l1` to use weighted-force directed algorithm for network layout. `-B1` to use white color background.
4. You should get a view similar to Fig. 10 (please see Supplementary Video 2 for a rotating 3D animation).

**Figure 10.**
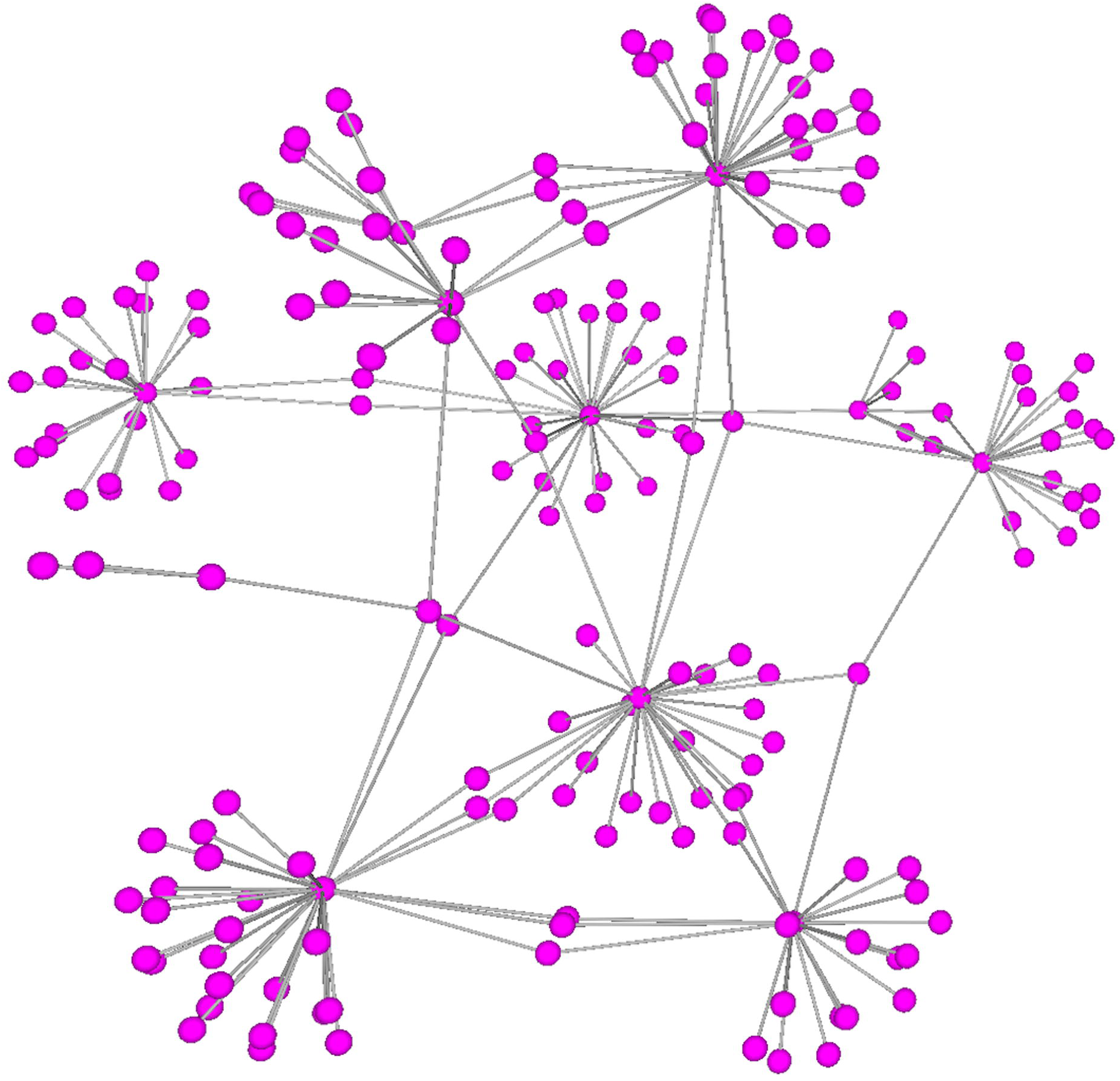
View of the network in weighted-force directed layout.

#### In-position visualization

In this approach, the current network layout (e.g. weighted force directed, simulated annealing force directed, linlog, etc.) stays the same post-clustering. As the positions of nodes are static, clustering information of nodes is communicated through node coloring. This approach is especially helpful when used in combination with layout algorithms that naturally position highly connected nodes closer (e.g. weighted force directed, and simulated annealing force directed).

5. In main menu, select a clustering algorithm from “Clustering Algorithms —>” list options (e.g. select “Clustering Algorithms —>Modularity” for modularity clustering algorithm).
6. You should get a view similar to as in Fig. 11.

**Figure 11.**
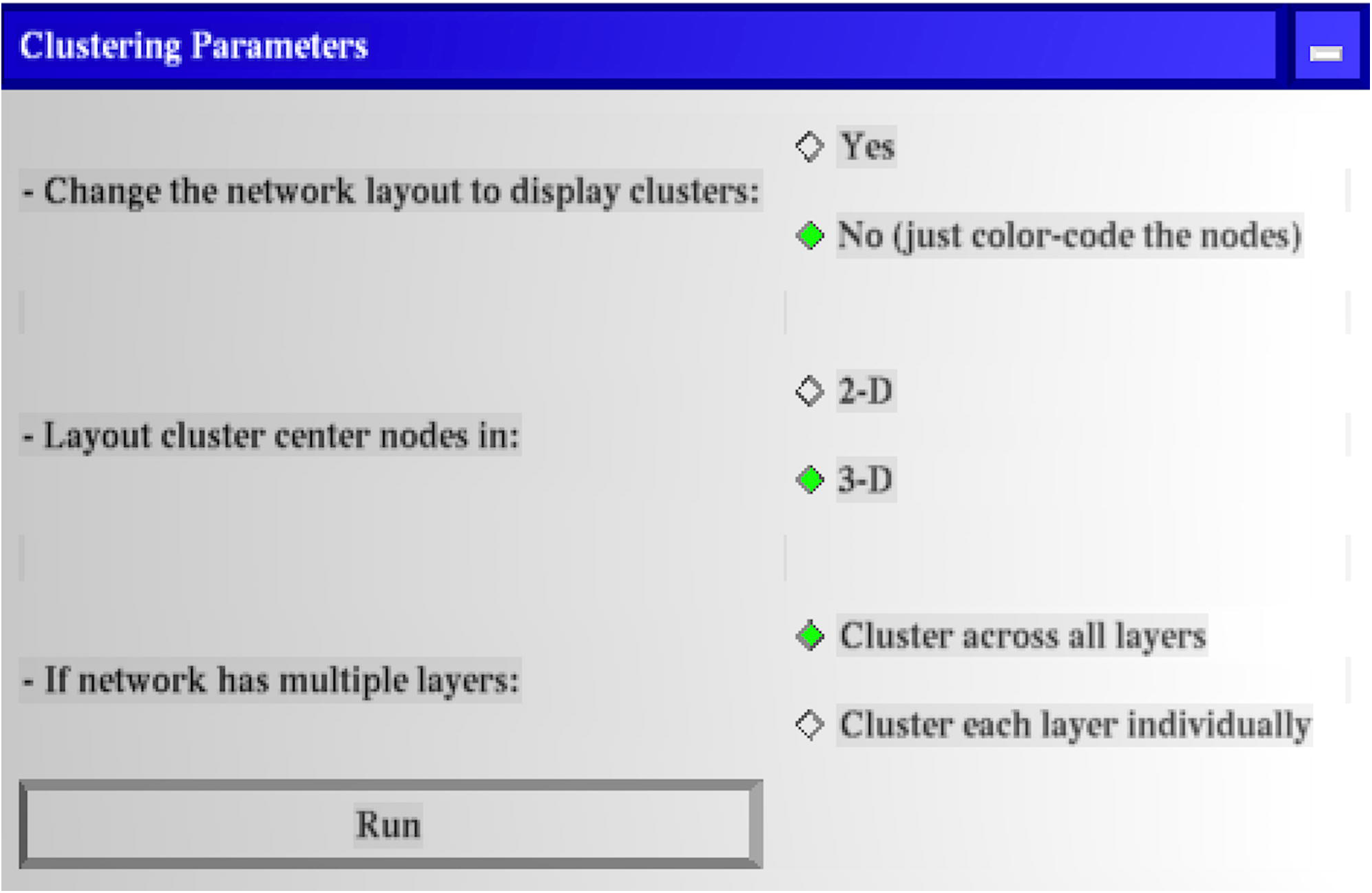
Clustering operation dialog box.
7. In this dialog box, choose the settings as shown in Fig. 11.
8. Click the Run button.
9. You should get a view similar to those in Fig. 12a-c. Each panel displays the same network layout with node colors corresponding to different clusters from different clustering algorithm results. These clusters visually overlap quite well with the weighted force directed algorithm used for the network layout.

**Figure 12.**
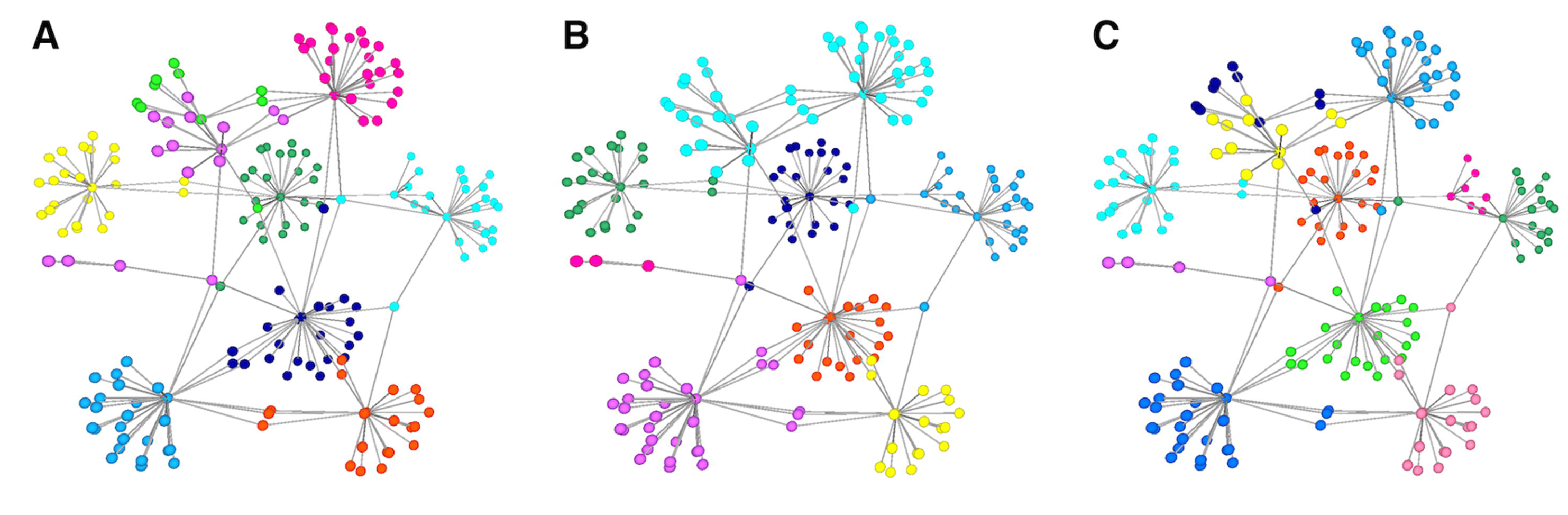
In-position visualization layout of the network using (a) modularity clustering, (b) Markov clustering, (c) edge-betweenness clustering algorithm.

#### 3D Circos layout visualization

3D Circos layout is another cluster visualization algorithm. Here, nodes are positioned on the surface of a 3D hemisphere sliced into pie-like panels that correspond to each cluster.

10. After Step 9, from the main menu select “Network Algorithms —> Create 3D Circos” button.
11. Assuming that you selected modularity clustering algorithm in step 5, you should get a network view similar to Fig. 13.a. Each cluster, in a different color, is on the surface of the 3D hemisphere in a pie-like panel.

**Figure 13.**
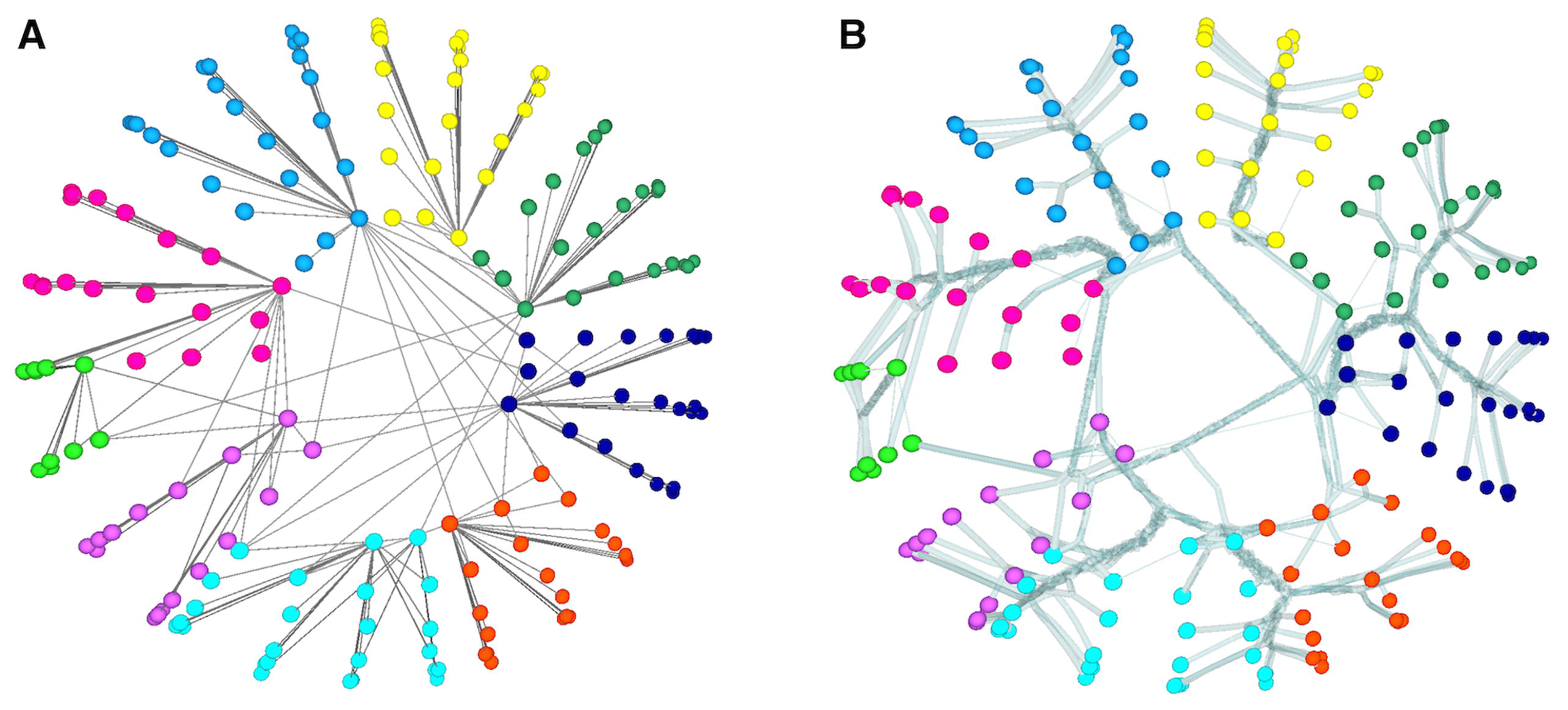
3D Circos layout of the network using modularity clustering with (a) no bundling, (b) edge bundling.

#### (optional) Edge Bundling

Optionally, the network in Fig. 13.a. can be further refined with edge bundling to visually clarify the strength of intra-cluster and inter-cluster connectivities. For example, when there are many edges between two clusters, the bundled edges are thicker, representing stronger connectivity. Reverse is true if there are only few edges between two clusters. The protocol steps for edge bundling are:

1. From the main menu, select “Network Algorithms —> Bundle Edges” button.
2. In the edge bundling dialog box, change the option for “Coloring of bundled edges” to “Use single bundling color”.
3. Click the Run button.
4. You should get a view similar to Fig. 13.b. with the modularity clustering algorithm. Similarly, Fig. 14 illustrates the network view following Markov (Fig. 14.a.) and edge-betweenness (Fig. 14.b.) clustering algorithms with edge bundling. Intra-cluster and inter-cluster connectivity strengths are much more clear. Because of edge bundling, overall connectivity among clusters can be discerned better compared to individual edges drawn as in Fig. 13.a. You can re-run the same operation by choosing different values for ECT and other parameters in the Edge Bundling dialog box and compare the network layouts.

**Figure 14.**
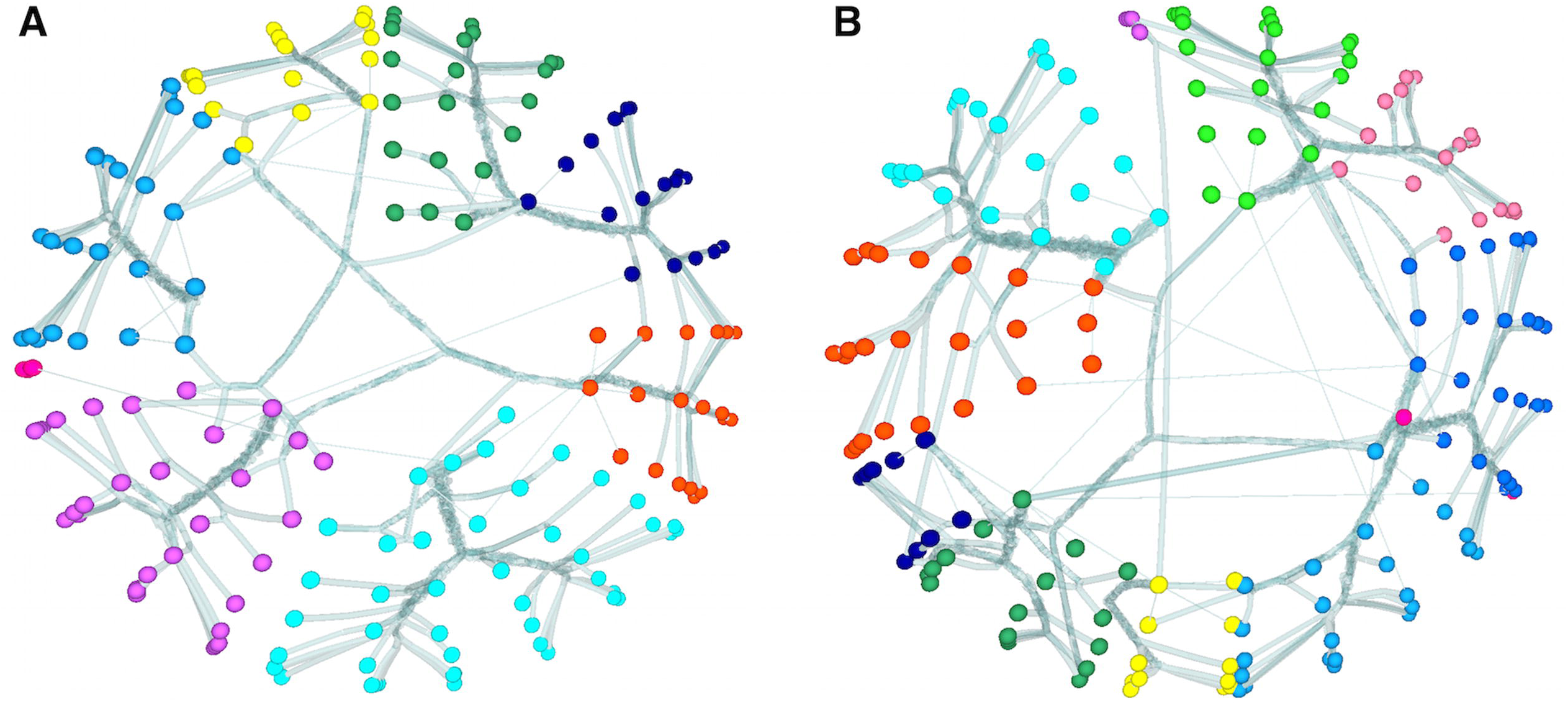
3D Circos layout of the network with edge bundling using (a) Markov clustering, (b) edge-betweenness clustering.

Please note that the edge bundling option is also useful in exploring the global structure of moderate-to-highly connected networks.

#### Force-directed layout visualization

Here, we illustrate when each cluster is positioned in-space after force-directed layout algorithm. Inside each cluster, nodes are organized using the hemispherical layout. As iCAVE users can select the background color in white or black, and for some network visualizations the black background provides better results, we will use a black background in this illustration.

16. Open a new terminal window and launch iCAVE with the following syntax:

~~~
vrnetview -Gvisual-cortex.txt -O10 -l1
~~~
17. From the main menu, select “Clustering Algorithms —> Modularity” button.
18. In this dialog box:

i. Keep the first toggle option “Yes”.
ii. Change the second toggle option to “2-D” (this option is useful for journal figures).
iii. Click the Run button. You should get a view similar to Fig. 15.a. Each hemisphere is colored differently, corresponding to a different cluster.

**Figure 15.**
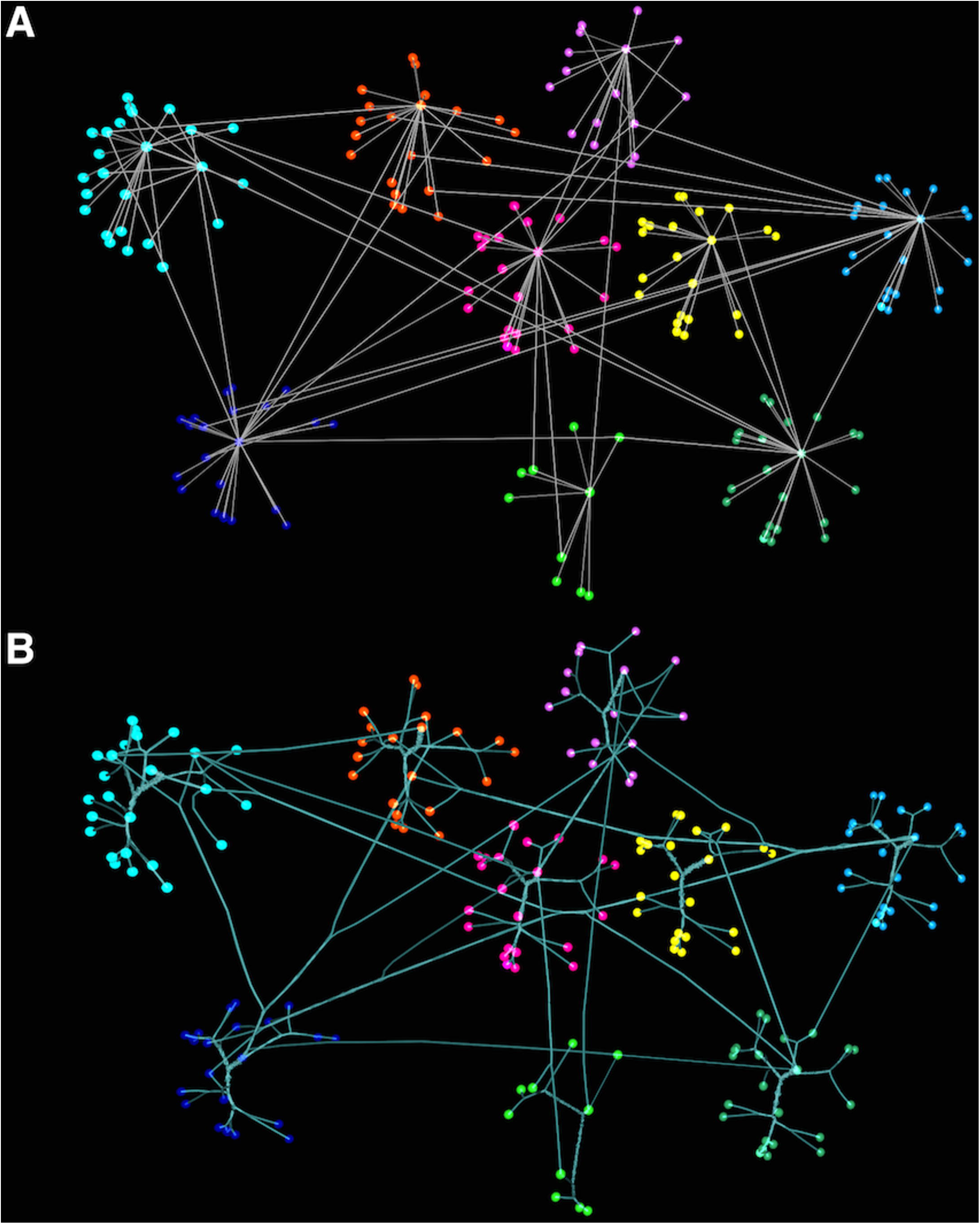
Force-directed layout of the network using modularity clustering with (a) no bundling, (b) edge bundling. To perform edge bundling:
19. From the main menu, select “Network Algorithms —> Bundle Edges” button.
20. In the edge bundling dialog box, change the following settings:

i. “Edge Compatibility Threshold (ECT)” value to 0.3.
ii. “Initial Step Size” value to 0.4.
iii. “Coloring of bundled edges” to “Use single bundling color”.
iv. “Blend the coloring of bundled edges” to “Yes”.
21. Click the Run button.
22. You should get a view similar to Fig. 15.b. Similarly, Fig. 16 illustrates the network view following Markov (Fig. 16.a.) and edge-betweenness (Fig. 16.b.) clustering algorithms (step 17) with edge bundling. Intra-cluster and inter-cluster connectivity strengths are seen in a much clearer way. You can re-run the same operation by choosing different values for ECT and other parameters in the Edge Bundling dialog box and compare the network layouts.

**Figure 16.**
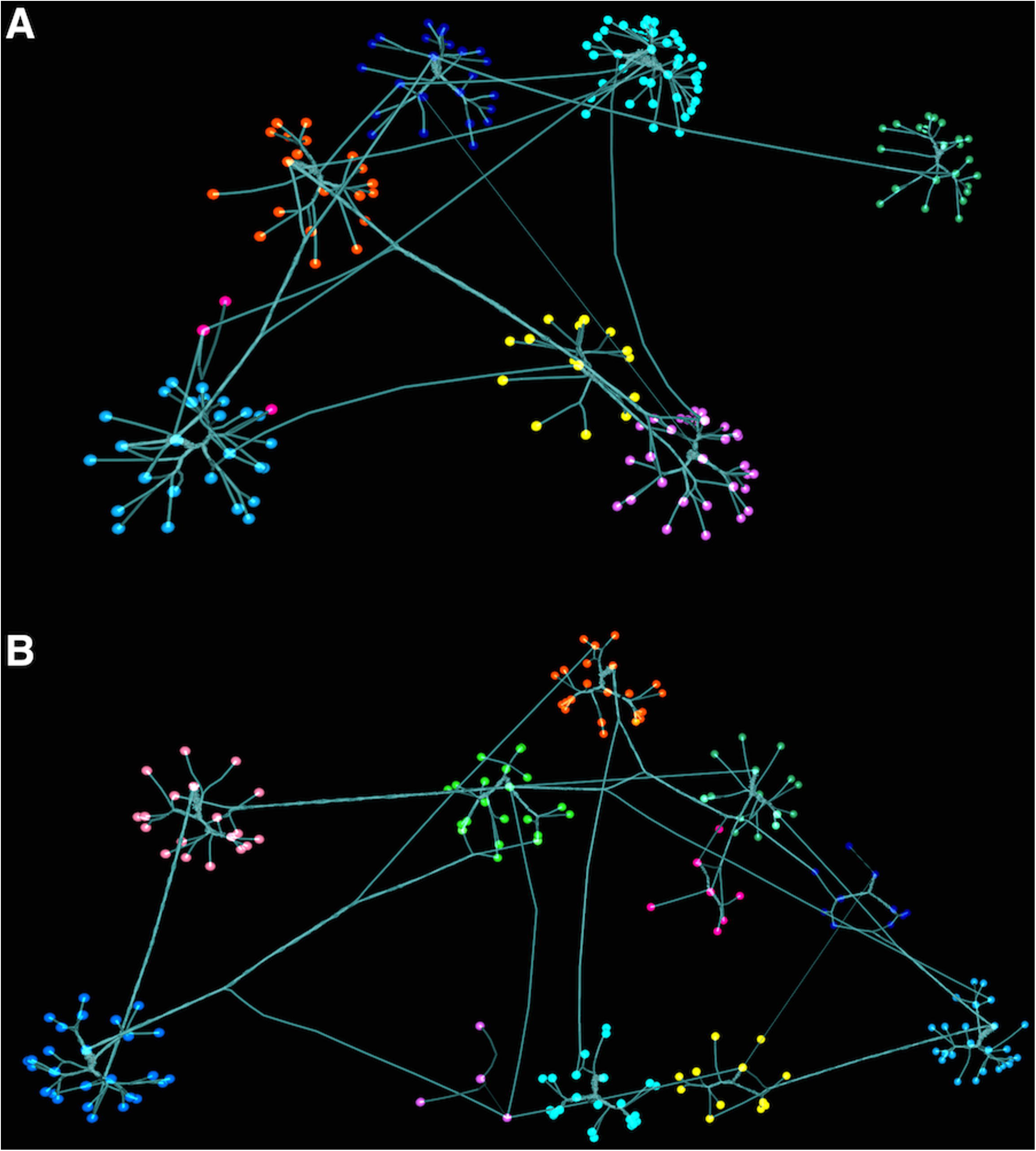
Force-directed layout of the network with edge bundling using (a) Markov clustering, (b) edge-betweenness clustering.

## BASIC PROTOCOL 4

### VISUALIZING NETWORKS WITH KNOWN 3D COORDINATES IN iCAVE

Basic Protocol 4 illustrates exploration of networks with known 3D coordinates. A wide-range of biological networks represent interactions between entities that have actual 3D physical coordinates, from residue-residue interaction networks within a protein to regional connectivity networks within a brain. These can be as simple as a plain 3D model of a biological structure, or coupled with additional layers of correlation information. It can also be extremely useful to visualize multiple networks within a known 3D biological structure (Doncheva, Klein, Domingues, & Albrecht, 2011), where visual comparisons between different networks within the same structure can provide clues on the underlying data patterns. Coupled with edge bundling, these visual explorations can help provide novel insights to better identify global network structures and generate hypotheses.

#### Necessary Resources

##### Input Files

We will first visualize residue-residue binary correlations within a G-protein coupled receptor (GPCR), which is a protein that traverses through the cell membrane, transmitting signals from extracellular to intracellular space. The correlation values are calculated from molecular dynamics (MD) simulation data of (Dror et al., 2013). In the input files, the edges represent pairwise correlations between the residues of a GPCR (M2 muscarinic acetylcholine receptor) for its three different states, to help better understand the effects of ligand binding to the protein.

The optional nodes section is the same for all three input files (i.e. “gpcr-wildtype.txt”, “gpcr-orthesteric.txt”, “gpcr-twoligands.txt”, see Internet Resources) and starts with:

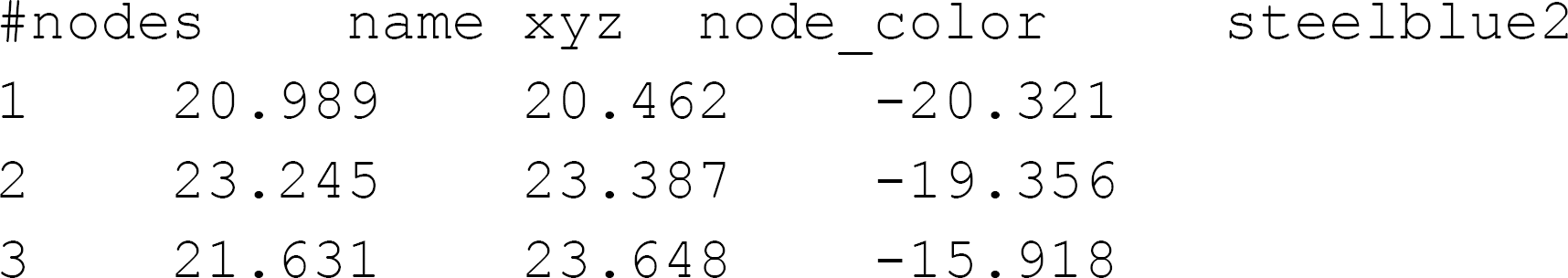

The header line includes the required ‘name’ attribute and optional ‘xyz’, and ‘node_color’ attributes. The optional ‘xyz’ attribute lets users specify x, y, and z coordinates of each node (as three tab-separated columns) in the input file. The optional ‘node_color’ attribute lets users specify a single color, ‘steelblue2’, for all the nodes in the network.

Protocol steps – *step annotations*

#### Protein Model

1. Copy input files “gpcr-wildtype.txt”, “gpcr-orthesteric.txt”, and “gpcr-twoligands.txt” into $iCAVE_HOME/vrnet/importFiles folder.
2. Open a terminal and change your directory to $iCAVE_HOME/vrnet.
3. Launch iCAVE with the following syntax:

~~~
vrnetview -Ggpcr-wildtype.txt -O10 -B1
~~~ `-Ggpcr-wildtype.txt` to open the graph specified in file gpcr-wildtype.txt. `-O10` to specify that the file is in iCAVE format. `-B1` to use white color background.
4. From the main menu, select “Adjust Node/Edge Size” toggle button.
5. Use the graphical sliders to adjust the size of nodes and edges.
6. Close the “Size Options” popup menu window by clicking on the top-right close button.
7. From the main menu of iCAVE, select “Network Algorithms —> Bundle Edges” button.
8. In the dialog box, use the graphical sliders and change

i. “Edge Compatibility Threshold (ECT)” value to 0.7.
ii. “Initial Step Size” value to 0.1.
9. Click the Run button.
10. You should get the network view similar to Fig. 17. a.

**Figure 17.**
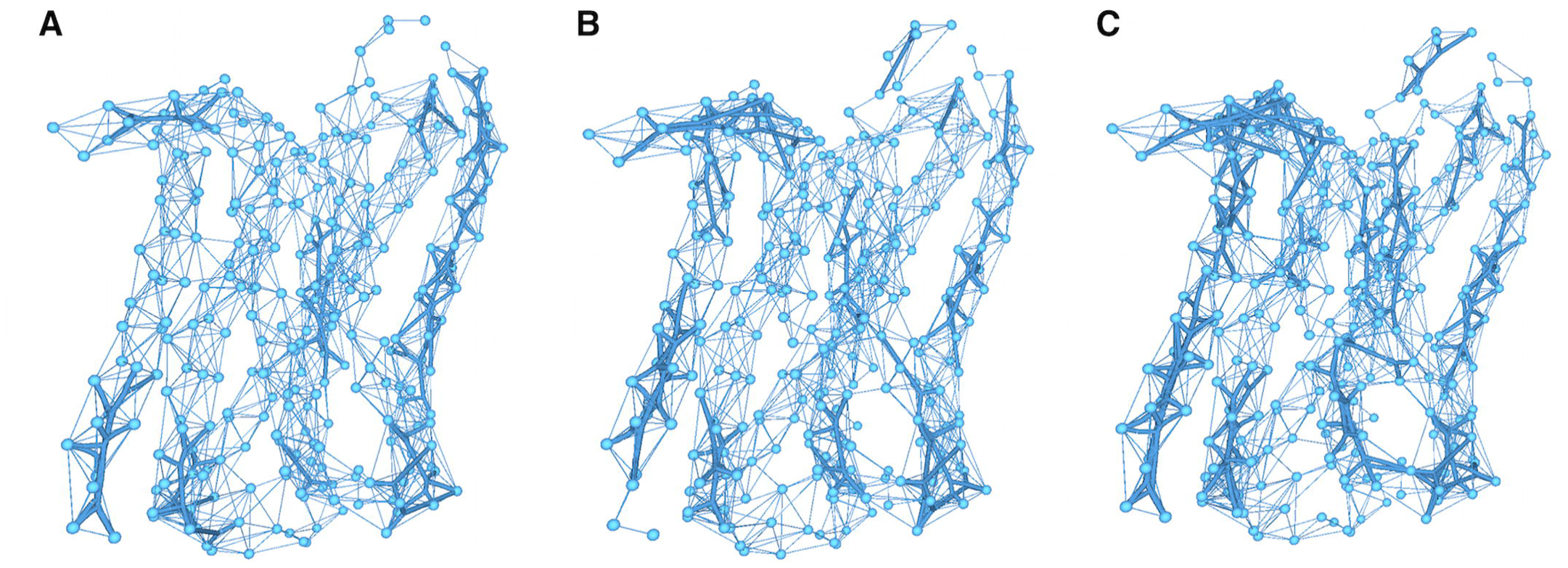
Edge-bundled view of the GPCR network in (a) wild type state, (b) orthesteric ligand-bound state, (c) both orthesteric and allosteric ligand-bound state.
11. Open a new terminal window and change your directory to $iCAVE_HOME/vrnet.
12. Launch iCAVE with the following syntax:

~~~
vrnetview -Ggpcr-orthesteric.txt -O10 -B1
~~~
13. Repeat steps 4-9.
14. You should get the network view similar to Fig. 17. b.
15. Open a new terminal window and change your directory to $iCAVE_HOME/vrnet.
16. Launch iCAVE with the following syntax:

~~~
vrnetview -Ggpcr-twoligands.txt -O10 -B1
~~~
17. Repeat steps 4-9.
18. You should get the network view similar to Fig. 17. c.

Comparison of these side-by-side figures reveals that ligand binding to the GPCR increases correlations between residue pairs.

Next, we provide examples on visualizing a brain connectome and a neuron. Both input files (“brain.txt”; “neuron.txt”, see Internet Resources) have formats similar to the protein network input files.

#### Brain Model

Brain connectome networks are widely utilized both for research and clinical purposes, and there are a large number of repositories that house such networks. For this protocol, we selected a human brain connectome generated by the PIT Bioinformatics Group (“The Budapest Reference Connectome Server v2.0,” 2015) using MRI scans obtained from the Human Connectome Project (McNab et al., 2013).

19. Copy input file “brain.txt” into $iCAVE_HOME/vrnet/importFiles folder.
20. Open a terminal and change your directory to $iCAVE_HOME/vrnet.
21. Launch iCAVE with the following syntax:

~~~
vrnetview -Gbrain.txt -O10 -B1
~~~ `-Gbrain.txt` to open the graph specified in file brain.txt. `-O10` the file is in iCAVE format. `-B1` to use white color background.
22. From the main iCAVE menu, select “Adjust Node/Edge Size” toggle button.
23. Use the graphical sliders to adjust the size of nodes and edges.
24. Close the “Size Options” popup menu window.
25. You should get the network view similar to Fig. 18.

**Figure 18.**
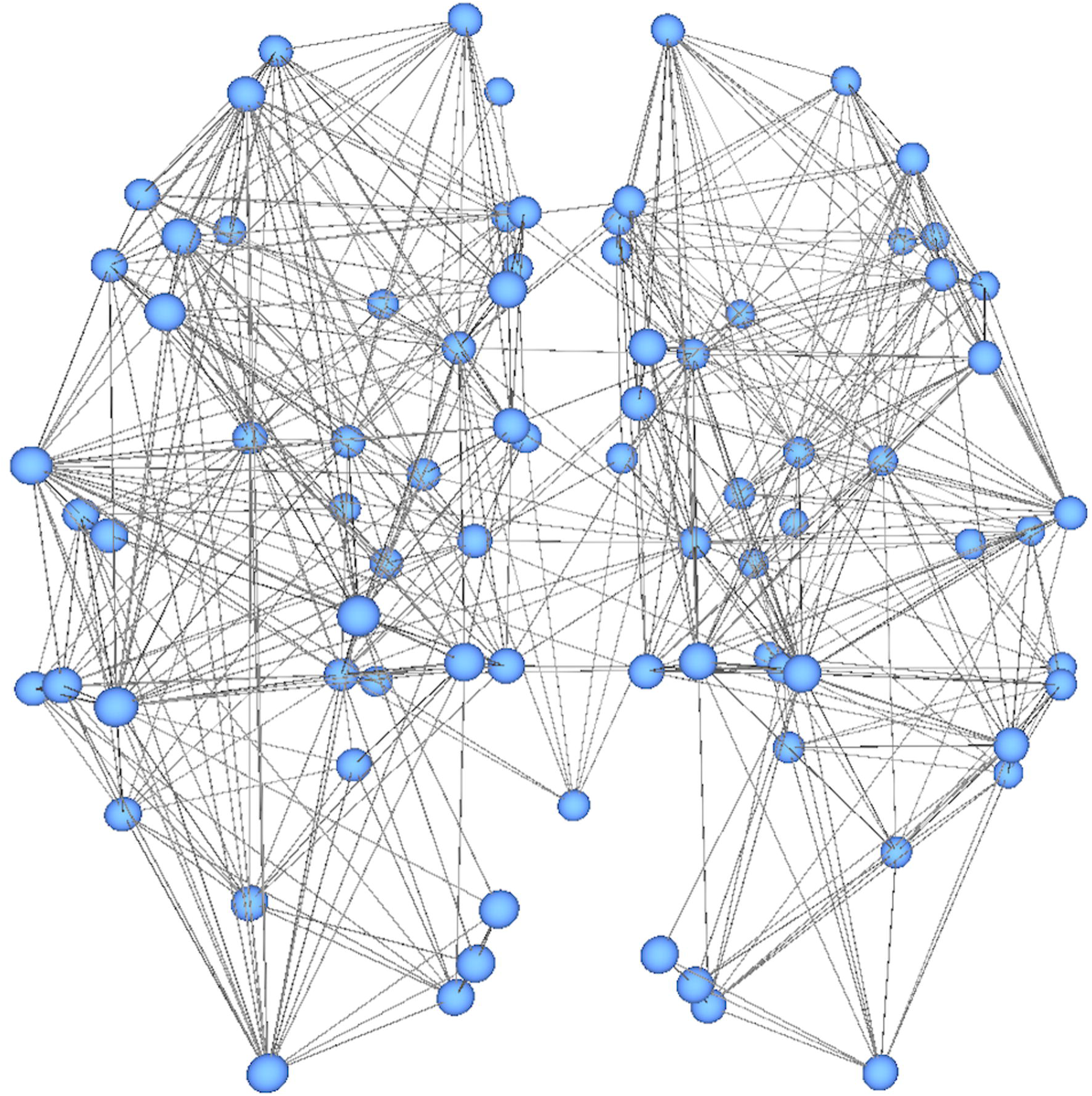
View of the brain connectome network.

#### Neuron Model

For this model, we first selected network data of human neuron morphology (Varga, Tamas, Barzo, Olah, & Somogyi, 2015) from NeuroMorpho.Org (Ascoli, Donohue, & Halavi, 2007), which provides a large collection of publicly accessible 3D neuronal reconstructions data for the neuroscience community. We then converted the file into iCAVE input format.

26. Copy input file “neuron.txt” into $iCAVE_HOME/vrnet/importFiles folder.
27. Open a terminal and change your directory to $iCAVE_HOME/vrnet.
28. Launch iCAVE with the following syntax:

~~~
vrnetview -Gneuron.txt -O10 -B1
~~~ `-Gneuron.txt` to open the graph specified in file neuron.txt. `-O10` the file is in iCAVE format. `-B1` to use white color background.
29. From the main menu, select and activate the “Show Single Nodes” toggle button.
30. From the main menu, select “Adjust Node/Edge Size” toggle button.
31. Use the graphical sliders to adjust the size of nodes and edges.
32. Close the “Size Options” popup menu window.
33. You should get the network view similar to Fig. 19.

**Figure 19.**
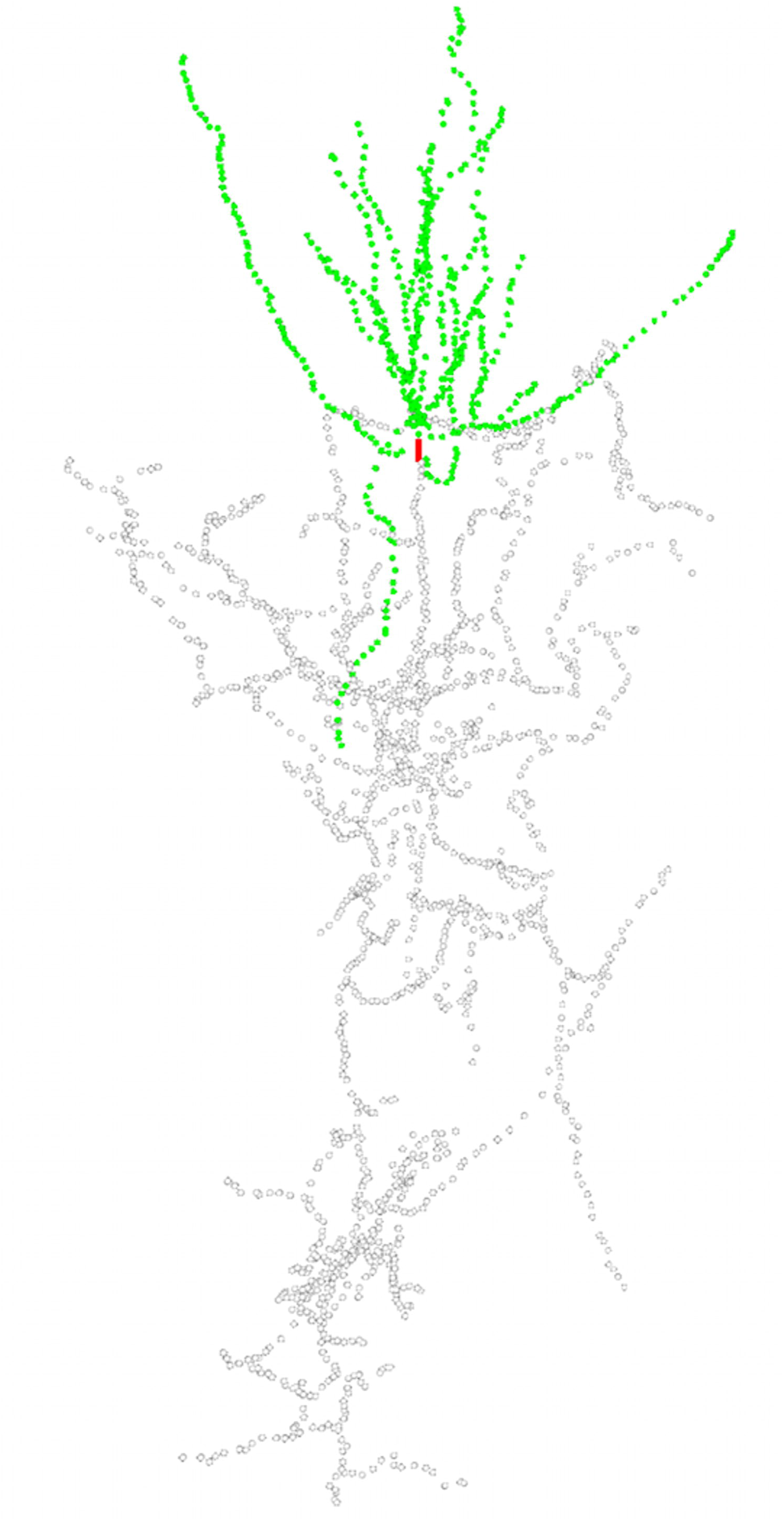
View of the human neuron reconstruction model.

## SUPPORT PROTOCOL 1

### INSTALLING iCAVE

iCAVE was developed using C++ and uses Vrui library based on OpenGL API platform. Before installing iCAVE, we require successful prior installation of Vrui and the libraries listed below.

#### Necessary Resources

##### Software

Operating System (OS): Linux, UNIX, MacOS

MacOS additional software:

#### iCAVE requires that XCode, XQuartz, libjpeg, libz, and libpng are already installed before installing iCAVE

Protocol steps:

1. Download the latest version of iCAVE from http://research.mssm.edu/gumuslab/software.html
2. Open a terminal window.
3. Change your working directory to the location where you saved iCAVE.
4. Open up the compressed package:

~~~
tar -xvzf $iCAVE PACKAGE NAME
~~~
5. Change your working directory to $iCAVE_HOME.
6. Start the installation process by running:

~~~
./install.sh
~~~
7. If the installation process finishes successfully, proceed to step 8. If any errors, please check our website for possible solutions.
8. After the successful installation of iCAVE, post-installation configuration tasks need to be performed. To achieve this, run:

~~~
source setup.bash
~~~
9. To test the iCAVE installation:

i. Open your X Window terminal.
ii. Make sure the necessary configurations are performed in your current terminal by running:

~~~
source setup.bash
~~~
iii. Change your working directory to $iCAVE_HOME/vrnet
iv. Launch iCAVE with a sample network input file with (sample file comes in the tar package):

~~~
vrnetview -Gsample.txt -O10
~~~
10. If you get an X Window terminal with the network visualization, you can start interactive explorations of the sample network with iCAVE. If you get any error message(s), please check the Troubleshooting section for possible solutions.

## SUPPORT PROTOCOL 2

### NECESSARY RESOURCES FOR STEREOSCOPIC 3D OPTION AND HOW TO TURN IT ON OR OFF

iCAVE is completely portable between 3D, stereoscopic 3D and immersive 3D environments. In this protocol, we describe the necessary hardware and software resources for 3D stereoscopic viewing, and the steps on how to turn on or off the stereo option.

#### Necessary Resources

Users can explore stereoscopic 3D visualizations using iCAVE if they have stereo-enabled hardware and active shutter glasses that convey stereoscopic images as described below. These glasses are significantly cheaper than recent head-mounted displays (HMDs). Alternatively, users can connect their stereo-enabled computers to large screens or curved walls, which provide the additional benefit of magnification and wide field of view, or interact with their networks in immersive CAVE environments. However, we would like to note that even without any of these resources, users can use iCAVE to interactively explore networks in 3D using their personal computers and utilize most its functionalities.

##### Hardware

*Stereo-enabled graphics card:* Stereoscopic 3D visualization on a 2D computer screen involves the generation of slightly different images for each eye, creating the perception of depth for the viewer. iCAVE supports professional-level quad buffered stereo which most modern computers come equipped with. If not, the computer can be upgraded with a compatible graphics card (e.g. Nvidia Quadro).

*Active shutter glasses:* Professional-level (active) stereo 3D viewer technology employs active shutter systems, which operate by presenting the image intended for the left eye while blocking the right eye, then presenting the right-eye image while blocking the left eye. This operation is repeated very rapidly so that the fusion of the two images is perceived as a single 3D image. Most modern active shutter 3D systems use liquid crystal (LC) shutter glasses (e.g. Nvidia 3D Vision) that are available at a sub-price range.

*120 Hz Monitor:* Due to the rapid switch between left-right eye view required by the active shutter systems, we recommend a monitor that can display at high refresh rates to prevent flickers, with a refresh rate of at least 100 Hz. Most modern LCD monitors support this refresh rate.

##### Software

Operating System: iCAVE runs in Mac and Linux systems for all its 3D functionalities. However, the stereo option requires a Linux-based operating system, as current Macs do not support quad-buffered stereo (yet).

Please note that iCAVE supports more basic (less restrictive; but lower quality) anaglyph stereo 3D option as well.

Protocol steps

1. To turn on the stereo option:

i. Open a terminal window.
ii. Change your working directory to $iCAVE_HOME.
iii. Run the command:

~~~
sh stereo.sh
~~~
2. To turn off the stereo option:

i. Open a terminal window.
ii. Change your working directory to $iCAVE_HOME.
iii. Run the command:

~~~
sh no-stereo.sh
~~~

## SUPPORT PROTOCOL 3

### SPECIFICATION OF INPUT FILES IN iCAVE

iCAVE provides a wide variety of options to customize the nodes and edges to better visualize, explore and communicate networks in 3D. iCAVE users can follow the examples provided in the basic protocols here, which have utilized several of these keyword and color options, and then apply them to their own networks. We provide a complete list of keywords and their descriptions for specifying node attributes in Table 1 and for specifying edge attributes in Table 2. Furthermore, iCAVE allows great flexibility in specifying edge and node colors, which are proven quite important in highlighting important parts of a network. A wide range colors can be specified in iCAVE, based on standards available in R software (Ihaka & Gentleman, 1996), which we summarize in ‘$iCAVE_HOME/vrnet/ColorChart.pdf

**Table 1.** Specification and customization of node attributes

| Keyword | Description |
| --- | --- |
| *name (required)* | Provide the unique name/id to all nodes listed. |
| *label* | You can specify node labels different than the node name to be displayed. You can also specify labels for a subset of nodes (leave label column blank for nodes without labels). |
| *xyz* | x, y, z coordinates of node; can be real or integer values. |
| *node_color* | To specify a single, unified color for all the nodes. In the next tab, provide the name of the color. |
| *color_code* | To specify different colors for various nodes. Provide the name of color for each individual node. |
| *texture_type* | To specify texture, provide an integer value accordingly with the following: 0: dashed node, 1: polka dot, 2: triangles in the circle, 3: striped, 4: stars. |
| *cluster* | Specify an integer id to associate each node with a pre-described cluster. |
| *foldChange* | Specify this when you want to input gene expression values, where the node color reflects the increase or decrease in expression of the gene the node represents. |
| *node_scale* | To change the scale of the nodes relative to each other; you can specify any real number value. |
| *labels_only_scaled_nodes* | When used in combination with *node_scale*, only the labels of scaled nodes (the ones with a different value than 0) are displayed. |
| *layer_code* | To specify a multi-layered network, specify the layer (use integer value) of each node. |
| *category* | To differentiate between different classes of nodes. If using superpathway, you can use the following codes: 0: protein, 1: complex, 2: rna , 3: mirna , 4: family, 5: abstract. If visualizing metabolic network, you can use the following codes: 0: unclassified, 1: amino acid, 2: acyl_carnitiness, 3: diacyl-PC, 4: lyso-PC, 5: acyl-alkyl-PC, 6: sphingomyeline. |
| *showNodeProperty* | If no *node_scale* option is provided and you specify this, each node is highlighted at intensity in accordance with its number of neighbors. |

**Table 2.**
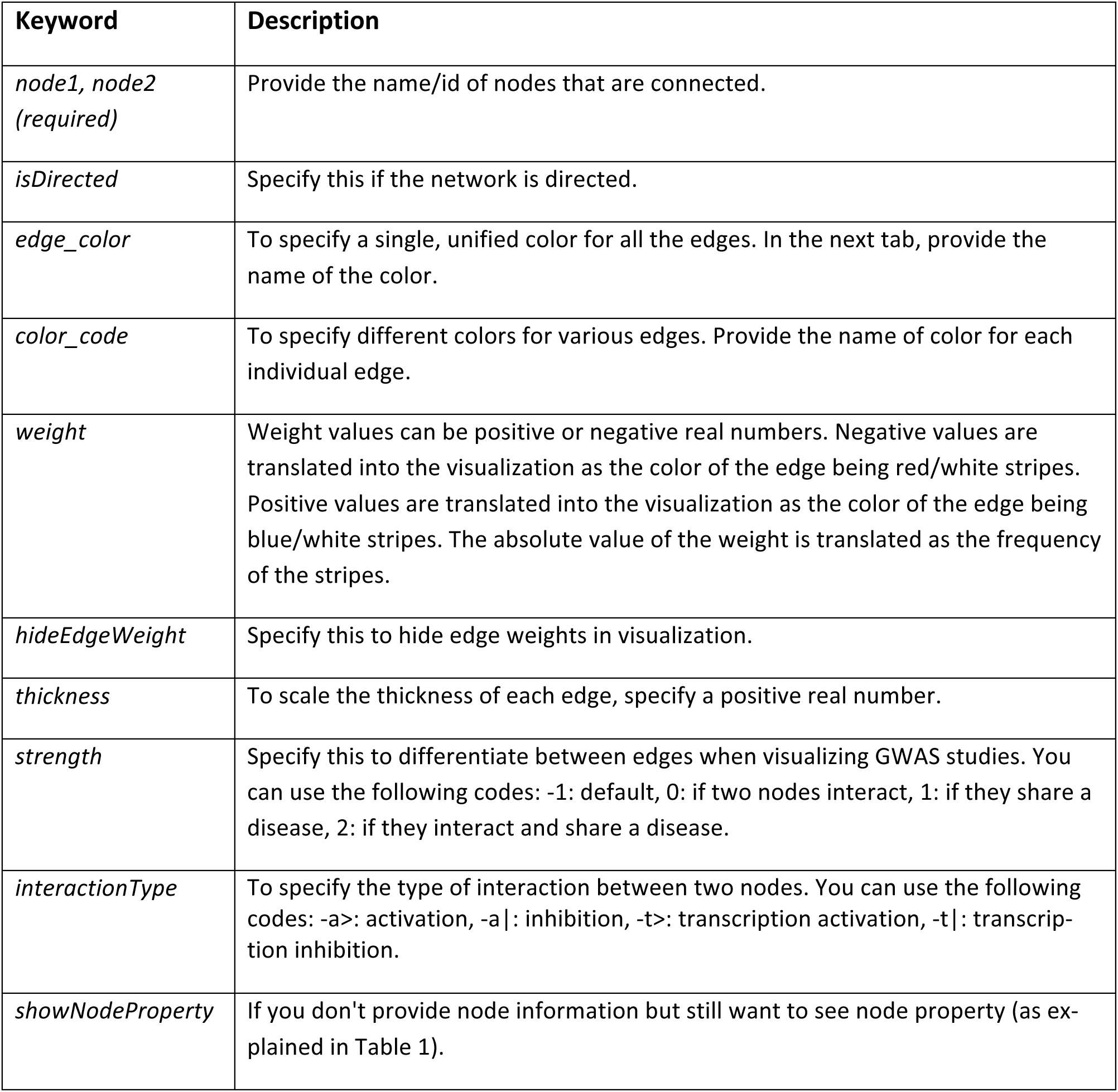
Specification and customization of edge attributes

### GUIDELINES FOR UNDERSTANDING RESULTS

To objective of iCAVE is to assist researchers in visualizing their networks to identify the unique characteristics of data that underlie large, dense or complex biological networks in a simple and user-intuitive fashion. Visualization of abstract networks in 3D is a nascent field, and how best to utilize features specific to 3D or to take advantage of the recent 3D technologies is currently an open research question. iCAVE can therefore also assist researchers in visualization design to perform user studies in order to better understand the relative advantages of its different 3D features. The protocols described here demonstrate different features of iCAVE for interactively exploring network visualizations in 3D, stereoscopic 3D and immersive 3D. To identify unique characteristics of the data that underlie large, dense or complex biological networks, researchers can build up on these basic protocols and utilize the full range of customization options in Support Protocol 3 to visually explore their networks. Overall, iCAVE is arguably the first 3D, stereoscopic 3D, and immersive 3D biomolecular network visualization tool that is open source and utilizable with commercial hardware/software and the responses and requests from its users will shape its future versions and in identifying its ideal usage scenarios.

### COMMENTARY

#### Background Information

Advances in computational and experimental technologies in life sciences lead to the generation of increasingly large and complex data, making it a real challenge to identify special patterns buried within them. Visually exploring the data in the form of network representations is often more user-intuitive than exploring the raw data. However, these networks are also increasing in size and complexity, and often contain multiple subtypes. For successful explorations, effective visualization tools that can accommodate such networks are needed.

We have recently developed iCAVE, an open-source network visualization tool to help researchers interactively explore large, complex, and multi-layered biological networks in 3D. In addition, iCAVE takes advantage of recent advances in 3D visualization technologies to enable users to visualize their networks in stereoscopic 3D (using an appropriate graphics card and stereo glasses) or immersive 3D (in a CAVE environment). The 3D environment provides additional navigational capabilities that let users view a complex network from multiple perspectives, and identify structures within the data that are difficult to discern in 2D. Being able to utilize the third dimension also lets users explore multi-layered networks with greater flexibility and explore networks with known physical 3D coordinates in their original spatial structure.

The network exploration and interaction functionalities within iCAVE 3D graphical user interface offers several alternative network layouts, clustering algorithms, and cluster visualizations to provide flexibility in addressing different user needs. Multi-layered networks or networks with physical 3D constraints can be visualized accurately. In addition, edge bundling functionality can be used during various exploration stages to identify global patterns or unique modules. iCAVE provides users with plenty of additional options to customize their networks, such as color, scaling, directionality, as summarized in Support Protocol 3.

Based on the continued interest from the research community in utilizing iCAVE for various biological problems, we are continuously enhancing its capabilities. Therefore, we accompany this protocol with an updated version of iCAVE, which includes these additional functionalities:

- Addition of ‘label’ node attribute. Users can now specify different labels for all or a subset of the nodes in their network. This feature is especially helpful when highlighting a small number of important nodes within a complex network.
- Clustering dialog box, which enables users to easily choose a cluster layout option.
- Edge bundling dialog box, for users to adjust edge bundling parameters (e.g. granularity) and specify different options for the visualization of the bundles (e.g. coloring or color blending).

The Basic Protocols provided here demonstrate some of the most common usage scenarios, though many other scenarios are possible. In general, network exploration is an iterative process, and accordingly, lessons learned at various steps can be pieced together strategically to improve the exploration experience and success.

#### Critical Parameters and Troubleshooting

##### Unable to load network

*Symptom 1:* When loading a network from an input file, you get an error message and iCAVE aborts launch process as it displays an error message similar to the following:

~~~
libc++abi.dylib: terminating with uncaught exception of type IO::File::OpenError: IO::StandardFile: Unable to open file importFiles/fileXYZ for reading due to error 2
~~~

*Possible causes:* iCAVE cannot locate the file that was passed as an argument.

*Remedies:* Check the location and name of the file in your machine and make sure it matches the argument passed to iCAVE.

*Symptom 2:* When loading a network from an input file, you get an error message and iCAVE aborts launch process as it displays an error message similar to the following:

~~~
edgecolor does not match to any known properties. Check the syntax and users manual
~~~

*Possible causes:* Input file was not generated accordingly with the iCAVE format specifications. For instance, wrong keyword was used.

*Remedies:* Check the error message and make sure your input file complies with iCAVE format specifications. For instance, the error message above complains about the usage of ‘edgecolor’ keyword in the input file. The problem can be resolved by changing it to the proper keyword, which is ‘edge_color’.

##### Takes long time to load network

*Symptoms:* When loading a network from an input file, it takes a long time to launch iCAVE graphical interface and display the initial network view without any error messages.

*Possible causes:* Network consists of large number of elements (nodes, edges) and/or the choice of layout algorithm may be taking a long time. Also, it is possible that the optional annotations (e.g. labels) in the input file increase the processing time.

*Remedies:* Change the network layout option used during the launch of iCAVE. Some layout algorithms take longer time than others to calculate positioning of nodes, which may delay the loading process. Try launching iCAVE with a less compute-intensive layout algorithm, such as hemisphere layout (argument -l2), or semantic levels (argument -l3) algorithm.

If the ‘label’ keyword is not used in the network specification file, iCAVE uses ‘name’ of all the nodes listed in the input file to use as node labels. During launch time, iCAVE generates all the necessary labels and makes them ready to use. This process can take a while if a large number of labels need to be generated. To speed up this process (and launch time), try to limit the number of node labels specified for your network using the ‘label’ keyword in accordance with iCAVE formatting guidelines.

##### Takes long time to perform an operation

*Symptoms:* After an operation is requested in iCAVE, the (visual) results don't appear for a long time and the program seems frozen.

*Possible causes:* Requested operation requires too much computation time.

*Remedies:* Some functionalities provided by iCAVE require considerable amount of computation, especially for networks with large number of nodes and/or edges. In some cases, adjusting the choice of input parameters can reduce the computation time. For instance, edge bundling is a computationally intensive operation in networks with large number of edges. In the edge bundling dialog window, increasing the “Edge Compatibility Threshold (ECT)” parameter value and/or lowering the value of other parameters result in less computation time to complete the operation. If the results are not satisfactory, this can be repeated multiple times as necessary and parameter values can be fine-tuned to balance the quality of results and computation time.

## ACKNOWLEDGEMENT

The authors gratefully acknowledge start-up funds to ZHG from the Department of Genetics and Genomics and the Icahn School of Medicine at Mount Sinai, and National Institutes of Health grant under award number 5 U19 AI118610. We also thank Dr. Sezen Vatansever for generating the GPCR example input file in Basic Protocol 4.

## INTERNET RESOURCES

http://research.mssm.edu/gumuslab/icave/cpbi-files

*Network input files referred in this manuscript can be found here.*

## VIDEOS

**“diseasome.gif” - Supplementary Video 1** Rotating 3D animation of the diseasome network.

**“visual-cortex.gif” - Supplementary Video 2** Rotating 3D animation of the mouse visual-cortex network.

